# Inhibition of formate production blocks CD8^+^ T-cell responses and delays disease onset in a mouse model of type 1 diabetes

**DOI:** 10.1101/2025.07.04.663229

**Authors:** Gabriela Ramirez-Hernandez, Micah Bell, Byungsoo Kong, Samuel Block, Matthew G. Vander Heiden, Nora Kory

**Affiliations:** Department of Molecular Metabolism, Harvard T.H. Chan School of Public Health, Boston, MA, USA; Dana-Farber Cancer Institute, Boston, MA, USA; Broad Institute of MIT and Harvard, Cambridge, MA, USA; Department of Biology, MIT, Cambridge, MA, USA; David H. Koch Institute for Integrative Cancer Research, MIT, Cambridge, MA, USA

## Abstract

The one-carbon metabolic pathway is essential for proliferating cells and has recently been identified as an immunomodulatory target in CD4⁺ T cells. However, its role in other immune cell types has not been fully established. We investigated the function of the one-carbon pathway in CD8⁺ T cells, which are the primary effectors responsible for the destruction of pancreatic beta cells that causes type 1 diabetes. Enzymes involved in the one-carbon pathway, as well as levels of formate—a critical intermediate—were upregulated during CD8⁺ T-cell activation. Pharmacological inhibition of MTHFD2, a mitochondrial enzyme involved in one-carbon metabolism, suppressed CD8⁺ T-cell activation, proliferation, and effector function. Mechanistically, this effect was mediated by reduced signaling through KRAS and the mTORC1 downstream targets HIF1α, S6, and STAT3. As previously shown in CD4⁺ T cells, formate supplementation reversed the effects of MTHFD2 inhibition on activation, proliferation, and function of CD8^+^ T cells, and prevented the reduction of the TCF1^high^ CD8⁺ progenitor cell population, which has been shown to drive anti-beta cell autoimmunity. Formate levels were elevated in the immune cells isolated from pancreatic lymph nodes during the insulitis stage in non-obese diabetic mice. Treatment of euglycemic non-obese diabetic mice with an MTHFD2 inhibitor during the insulitis stage delayed CD8⁺ T-cell infiltration into pancreatic islets and postponed the onset of type 1 diabetes. These findings reveal a new paradigm for preventing and delaying the onset of type 1 diabetes.

## Introduction

Type 1 diabetes (T1D) is a multifactorial, heterogeneous disease caused by a complex interaction of genetic, immunological, and environmental factors. This complexity complicates prevention efforts and poses challenges for treatments beyond permanent insulin replacement. Currently, T1D incidence is increasing at alarming rates in both children and adults, especially following coronavirus infection (Debuysschere et al. 2024; Ogle et al. 2022), highlighting the urgent need for new and effective therapies (Herold et al. 2024).

A hallmark of T1D is the selective autoimmune destruction of insulin-producing pancreatic islet β cells. CD8^+^ T cells play a crucial role in T1D development, acting as the primary agents responsible for β-cell killing (Dwyer et al. 2021; Gearty et al. 2022). Therefore, therapeutic strategies have targeted these cells for treating T1D, although with limited success (broadly reviewed in Herold et al., 2024, and Yang et al., 2024). A more in-depth understanding of the mechanisms underlying autoreactive CD8^+^ T cells will enable the development and improvement of interventions for T1D and help bridge this gap.

The CD8^+^ T-cell response requires metabolic reprogramming to meet the increased bioenergetic and biosynthetic demands of proliferation and effector function maintenance. Transcriptome analysis of activated CD8^+^ T cells revealed that one-carbon (1C) metabolism is among the most induced metabolic pathways (Ron-Harel et al., 2016; Ma et al., 2017). 1C metabolism is crucial for rapid cell proliferation, providing 1C units essential for biosynthetic processes, chromatin regulation, amino acid homeostasis, and redox defense (Ducker and Rabinowitz 2017). Its role and therapeutic applications in various cancers, including breast cancer, lung cancer, leukemia, and melanoma, have been extensively studied (Ducker and Rabinowitz 2017; Ren et al. 2024). In parallel, combination immunotherapies have emerged as a new paradigm in cancer treatment, prompting investigations into the role of 1C metabolism in immune cells (Ren et al. 2024). However, targeting this pathway for autoimmune disease remains largely unexplored.

The antifolate drug methotrexate (MTX), which inhibits dihydrofolate reductase (DHFR) in 1C metabolism, is a mainstay in the treatment of some autoimmune disorders, including rheumatoid arthritis and lupus. However, its benefits in T1D are not clear (Buckingham and Sandborg 2000), and it is not used clinically for this indication. Moreover, MTX toxicity extends to many proliferating tissues due to the broad expression of DHFR in all dividing cells (Ducker and Rabinowitz 2017; Yujing Li et al. 2025). Therefore, more selective inhibition of immune cell 1C metabolism may offer clinical benefits with reduced off-target effects.

Recent work identified methylenetetrahydrofolate dehydrogenase 2 (MTHFD2), a mitochondrial 1C pathway enzyme, as a control point of CD4^+^ T-cell differentiation and function. This suggests an anti-inflammatory role for inhibitors of this enzyme and implicates it as a promising target for autoimmune diseases such as multiple sclerosis (Sugiura et al. 2022). Additionally, MTHFD2 is highly expressed in immune and cancer cells, but is low or absent in normal adult tissues (Nilsson et al. 2014; Yujing Li et al. 2025). Whether MTHFD2 and its metabolic products influence the CD8^+^ T-cell response in an autoimmune diabetes context remains unknown. We investigated the role of MTHFD2 and the downstream metabolic product of the mitochondrial 1C pathway— formate— in the CD8^+^ T-cell response and assessed the potential of targeting this pathway in T1D. We identified that formate production is upregulated during CD8^+^ T cell activation associated with development of T1D in non-obese diabetic (NOD) mice. Pharmacological inhibition of MTHFD2 impaired CD8^+^ T-cell survival, proliferation, and function, via effects on signaling downstream of KRAS and mTORC1, and these effects were rescued by exogenous formate. Moreover, we found that inhibition of MTHFD2 delays T1D onset. Additionally, we found that MTHFD2 and formate modulate the expression of TCF1, a key transcription factor involved in T-cell memory, longevity, and self-renewal (Gearty et al., 2021). Together, these data demonstrate the impact of MTHFD2 and formate on CD8^+^ T-cell response and support their potential as therapeutic targets in autoimmune conditions.

## Results

### 1C metabolism enzymes and the metabolite formate are upregulated during early CD8^+^ Tcell response

After stimulation, T cells exit quiescence to initiate their expansion, effector function, and differentiation, one of the hallmarks of which is reprogramming of mitochondrial metabolism (Chapman, Boothby, and Chi 2020). Analysis of the mitochondrial proteome and RNA transcriptome of CD4^+^ and CD8^+^ T cells, respectively, identified 1C metabolism—particularly its mitochondrial branch—as the main induced metabolic pathway during early T-cell activation (effector function and expansion) (Ron-Harel et al. 2016; Ma et al. 2017).

To understand the dynamics of 1C metabolism during the CD8^+^ T-cell response, we mined a proteome dataset generated by ProteomeXchange (Gassaway et al. 2024) and analyzed the abundance of key proteins involved in central mitochondrial metabolic pathways during *in vitro* (anti-CD3 and anti-CD28 [aCD3/CD28] stimulation) induction of CD8^+^ T-cell activation. Proteome analysis identified 1C metabolism as a highly upregulated pathway, highlighting thymidylate synthase (TYMS) and MTHFD2 (Fig. 1A), enzymes essential for the biosynthesis of purines and thymidine. This was consistent with the reanalysis of the transcriptome database of *in vivo* CD8^+^ T-cell activation (infection with Lm-OVA) generated by the Immunological Genome Project transcriptome (Best et al. 2013; Ma et al. 2017) (Fig. S1), and with previous reports (Ron-Harel et al. 2016; Ma et al. 2017). Indeed, we observed an increase in 1C metabolism enzymes (Fig. 1B) during *in vitro* CD8^+^ T-cell activation by aCD3/CD28 with IL-2 (Fig. 1C).

**Figure 1:**
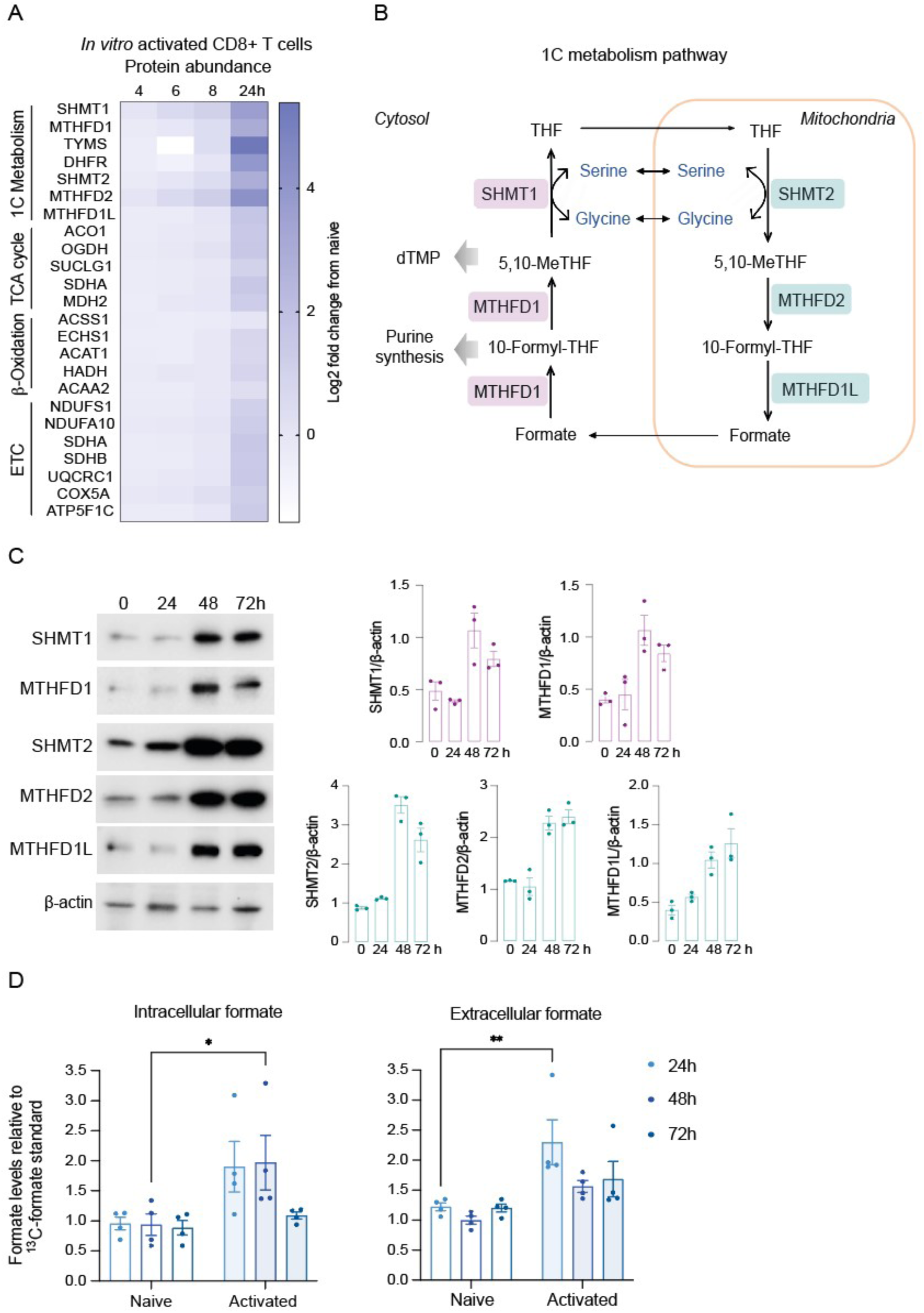
1C metabolism and formate production are induced during the CD8⁺ T-cell response. (A) Heatmap of proteomic datasets displaying the fold change of individual key proteins within the main mitochondrial metabolic pathways during the *in vitro* (aCD3/CD28 stimulation)–induced CD8⁺ T-cell effector response, compared with naïve cells (0h). (B) Schematic of 1C metabolic pathway with cytosolic and mitochondrial enzymes highlighted in purple and teal, respectively. (C) Immunoblot of the cytosolic (SHMT1, MTHFD1) and mitochondrial (SHMT2, MTHFD2, and MTHFD1L) branches of the 1C metabolic pathway enzymes during *in vitro*–activated CD8⁺ T cells (aCD3/CD28, IL-2 stimulation). Relative expression to b-actin. (D) Intracellular and extracellular formate levels per 2×10^6^ cells relative to standard sodium ^13^C-formate, measured by GC-MS in naïve and *in vitro*–activated CD8⁺ T cells. Data are shown as mean ± SEM. Statistical multiple comparisons performed by two-way ANOVA: *p < 0.05, **p < 0.01.

A central intermediate metabolite of 1C metabolism is mitochondrially produced formate, which is crucial for supporting the downstream functions of 1C metabolism. The sole carbon of formate is incorporated into purine and thymidylate synthesis, production of the amino acid serine from glycine, and can contribute to methylation reactions (Lamarre et al. 2013; Ducker and Rabinowitz 2017). Despite having a central role in cellular metabolism, formate levels during T-cell activation have not been studied.

To explore cellular formate levels, we adapted and applied a gas chromatography-mass spectrometry (GC-MS) method for detecting and quantifying formate base on a previously described protocol (Lamarre et al. 2014). This technique, capable of quantifying formate levels at low concentrations present in cellular environments, chemically derivatizes formate with 2,3,4,5,6-pentafluorobenzyl bromide (PFBBr) to produce a larger compound allowing detection by GC-MS (Fig. S2A). The resulting formate derivative, pentafluorobenzyl formate, exhibited a distinct and unique spectral peak and showed linearity up to a concentration of 500 µM (Fig. S2B).

Measurement of both intra- and extracellular formate levels during *in vitro* CD8^+^ T-cell activation showed a significant increase in formate production and secretion (Fig. 1D). Intracellular formate levels peaked at 48 hours post-activation, coinciding with the onset of T-cell proliferation and upregulation of the 1C metabolism enzymes responsible for formate production (Fig. 1C). In contrast, extracellular formate levels peaked earlier, at 24 hours post-activation, suggesting that in response to T-cell expansion, cells increase formate production while ceasing its secretion. A similar dynamic was observed in activated CD4^+^ T cells (Fig. S2C). Altogether, these findings highlight the changes within the 1C pathway and its metabolite formate during T-cell responses.

### Inhibiting the mitochondrial 1C metabolism enzyme MTHFD2 reduces formate production and hinders the CD8**⁺** T-cell response by suppressing KRAS and downstream targets of the mTOR pathway

An *in vivo* CRISPR-Cas9-based screen in primary CD4^+^ T cells identified MTHFD2 as an essential gene for T-cell proliferation and inflammatory function (Sugiura et al. 2022). This enzyme was also overexpressed in the whole blood of patients with various inflammatory and autoimmune diseases, including ulcerative colitis, Crohn’s disease, celiac disease, rheumatoid arthritis, systemic lupus erythematosus, and multiple sclerosis (Sugiura et al. 2022). Studies on the genetic deletion and pharmacological inhibition of MTHFD2 have established it as a control point for CD4^+^ T cells (Sugiura et al. 2022). Moreover, MTHFD2 suppressed the polarization of pro-inflammatory macrophages (M1) while enhancing the anti-inflammatory macrophage (M2) phenotype both *in vitro* and *in vivo* (Shang et al. 2023). While MTHFD2 has known strong effects on modulating other types of immune cells, including CD4^+^ T cells and macrophages, its role in CD8^+^ T cells remains unclear. Using the potent, selective, orally available, and recently discovered MTHFD2 inhibitor DS18561882 (Kawai et al. 2019), we evaluated the effect of inhibiting this enzyme on the CD8^+^ T-cell response stimulated by aCD3/CD28 with IL-2 *in vitro*.

The MTHFD2 inhibitor (MTHFD2i) reduced formate levels, preventing the accumulation previously observed 48 hours post-activation of CD8⁺ T cells (Fig. 2A). Moreover, viability (Fig. 2B), proliferation (Fig. 2C), and activation, as measured by CD44 expression (Fig. 2D), were decreased following MTHFD2 inhibition. In contrast, the inhibitory marker PD1 but not CD38, was increased (Fig. 2D). Additionally, the expression of IFNg, but not Perforin or Granzyme B, was significantly reduced, indicating a partially compromised effector function of CD8⁺ T cells (Fig. 2E). Thus, MTHFD2 is required for the optimal survival, expansion, activation, and function of CD8⁺ T cells *in vitro*.

**Figure 2:**
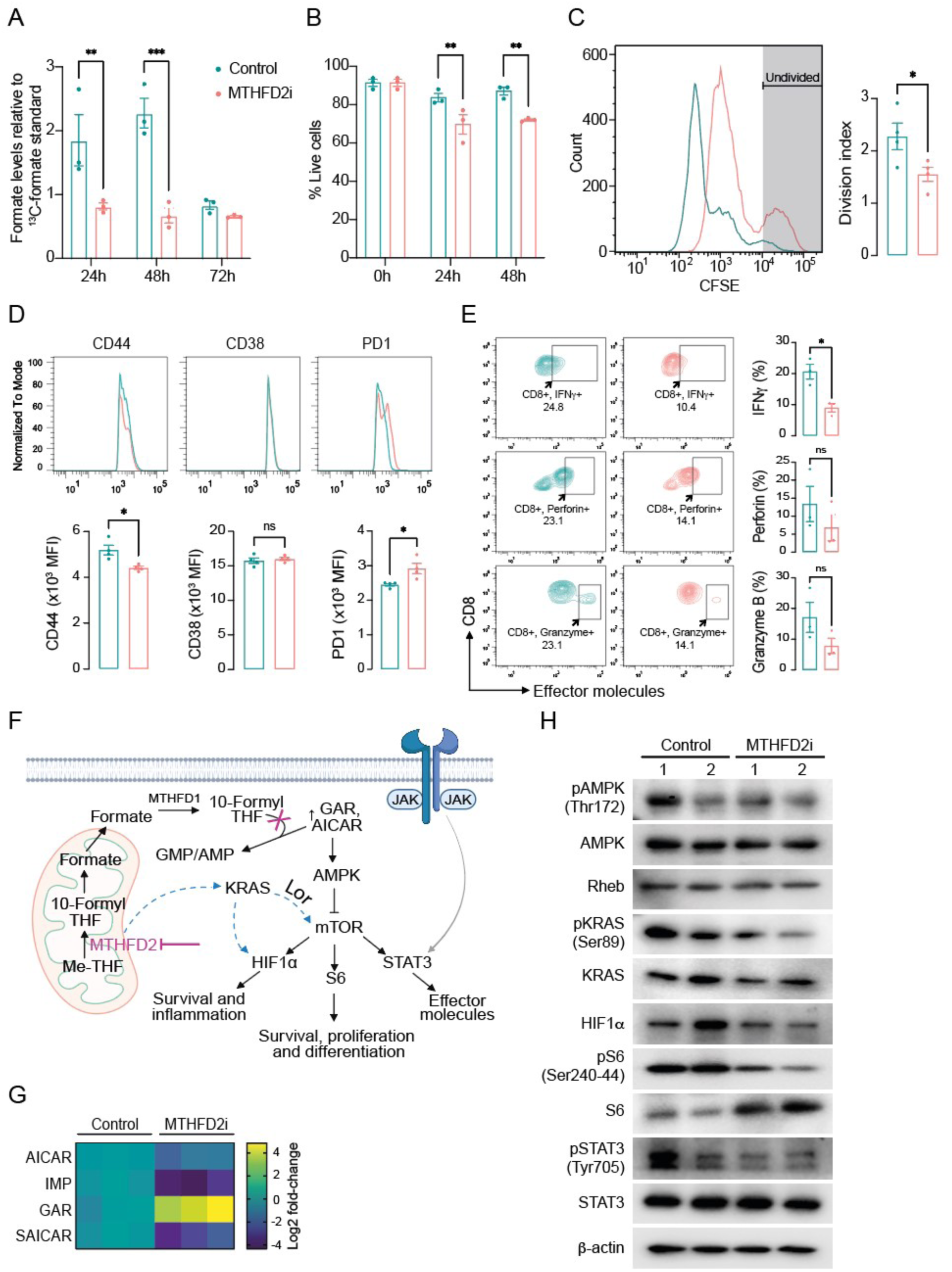
Inhibition of the mitochondrial 1C metabolism enzyme MTHFD2 and formate production impairs CD8⁺ T-cell survival, proliferation, activation, and function by suppressing KRAS and downstream targets of the mTOR pathway. (A) Intracellular formate levels relative to standard sodium ^13^C-formate, measured by GC-MS in 2×10^6^ activated CD8⁺ T cells treated with DMSO (control) or 500 nM MTHFD2 inhibitor (MTHFD2i) for 24, 48, and 72 hours. (B) Viability of activated CD8⁺ T cells treated with DMSO (control) or 500 nM MTHFD2i for 0, 24, and 48 hours. (C) Proliferation index measured by CellTrace CFSE dilution. (D) Flow cytometric analysis of the expression of activation marker CD44, inhibitory markers CD38 and PD1, and (E) effector molecules IFNg, Perforin, and Granzyme B in activated CD8+ T cells (gated on non-stimulated cells). (F) Schematic representation of the regulatory mechanisms of MTHFD2 in relation to the AMPK/mTOR pathway. MTHFD2 MTHFD2 influences AMPK/mTORC1 signaling by facilitating the accumulation of purine synthesis intermediates, but MTHFD2 can also act in other mTORC1 regulators, such as KRAS. (G) Heatmap of changes in purine synthesis intermediate abundance (AICAR, IMP, GAR, and SAICAR) measured by mass spectrometry. (H) Immunoblot of AMPK, phospho-AMPK, Rheb, KRAS, phospho-KRAS, HIF1α, S6, phospho-S6, STAT3, phospho-STAT3, and b-actin. Panels D, E, G, and H were generated using activated CD8⁺ T cells treated with DMSO (control) or 500 nM MTHFD2i for 48 hours; activation was induced by anti-CD3 and -CD28 (aCD3/CD28) antibodies and IL-2, and cells were re-stimulated with PMA and ionomycin for 4 hours before harvesting. Data are shown as mean ± SEM. Statistical multiple comparisons in panels A and B were performed by two-way ANOVA. Statistical comparisons between two groups in panels C-E were performed by unpaired t-test: *p < 0.05, **p < 0.01, ***p < 0.001, ns: non-significant.

To investigate the mechanism associated with the impaired CD8⁺ T-cell response under MTHFD2i conditions, we first examined the canonical enzymatic action of MTHFD2 (Fig. 2F). MTHFD2 supports cell proliferation and survival by providing key substrates for the synthesis of purine nucleotides through the dehydrogenation of 5,10-methylene tetrahydrofolate (5,10-CH₂-THF) and its conversion to 5,10-methenyl-THF, followed by subsequent ring opening to form 10-formylTHF, which is essential for *de novo* purine synthesis (Zarou, Vazquez, and Vignir Helgason 2021; Pardo-Lorente and Sdelci 2024; Liu et al. 2025). Moreover, levels of the purine synthesis intermediates GAR, SAICAR, and AICAR have been found to be elevated in MTHFD2-deficient HEK293T cells (Ducker et al. 2016) and during transient inhibition of MTHFD2 in primary Th1, Th17, and Treg cells (Sugiura et al. 2022).

We measured the levels of these metabolic intermediates in purine biosynthesis and noted an accumulation of the GAR intermediate with MTHFD2i treatment (Fig. 2G), which depends on 10formyl-THF to form FGAR during the third reaction in *de novo* purine biosynthesis (De Vitto et al. 2021), thus suggesting impaired purine synthesis.

The serine/threonine kinase mammalian target of rapamycin (mTOR) is one of the major regulators of T-cell activation, differentiation, and function (Pollizzi and Powell 2015; Saleiro and Platanias 2015). Additional roles of MTHFD2, beyond its mitochondrial enzymatic functions, have been reported, including an effect on the mTOR complex 1 (mTORC1) (Sugiura et al. 2022; Pardo-Lorente and Sdelci 2024).

To understand the mechanism(s) by which MTHFD2i controls CD8^+^ T-cell response, we assessed the mTOR downstream targets. We observed that the regulators of glycolytic metabolism, differentiation, and function of CD8⁺ T cells—HIF1α, S6, and STAT3—were repressed under MTHFD2i conditions, as indicated by HIF1α destabilization and S6 and STAT3 dephosphorylation (Fig. 2H).

Several reports have indicated that AICAR and guanine nucleotides activate AMPK and negatively regulate mTORC1 signaling in lung cancer, carcinoma, muscle, and T cells (Agarwal et al. 2015; Rao et al. 2016; Emmanuel et al. 2017; Zhan et al. 2018; Sugiura et al. 2022). However, we did not observe any changes in AICAR (Fig. 2G), AMPK, or Rheb levels (Fig. 2H). Akt is another critical regulator of mTORC1, and the Akt-mTORC1 signaling has been shown to be crucial for M2 polarization (Zhou et al. 2020). In the same manner as AMPK, Akt levels were not altered in the MTHFD2i conditions in CD8^+^ T cells (data not shown).

In colorectal cancer and pulmonary adenocarcinoma cells, activation of KRAS upregulates MTHFD2 (Yuchan Li et al. 2022; Pardo-Lorente and Sdelci 2024). Moreover, CD4⁺ and CD8⁺ T cells deficient in KRAS show significantly reduced activation, proliferation, and cytokine production in the context of graft-versus-host disease (Luo et al. 2020). We observed a reduction in KRAS activation during treatment with MTHFD2i (Fig. 2H), suggesting that in CD8^+^ T cells, MTHFD2 regulates the mTOR pathway through its effects on KRAS.

### MTHFD2 inhibitor effects on CD8^+^ T-cell response are mitigated by formate supplementation

Within the mitochondria, SHMT2, MTHFD2, and MTHFD1L facilitate the conversion of imported THF to formate for export and use in the cytosol (Green et al. 2023). Formate fulfills the 1C demand for the synthesis of nucleotides and methyl groups (Pietzke, Meiser, and Vazquez 2020), and formate supplementation (alone or in combination with other metabolites) can rescue T-cell survival, proliferation, and activation under serine deprivation or deletion of its mitochondrialproducing enzymes (SHMT2, MTHFD2) (Ron-Harel et al. 2016; Ma et al. 2017; Sugiura et al. 2022). To investigate these formate-rescuing actions, we treated activated CD8^+^ T cells under MTHFD2 inhibition conditions with or without exogenous formate. Formate levels measured by GC-MS indicated that formate supplementation elevated intracellular formate levels under control conditions and restored the diminished formate levels observed with MTHFD2 inhibition (Fig. 3A). As expected, CD8^+^ T-cell expansion began 48 hours after activation, and formate did not alter CD8^+^ T-cell proliferation; however, formate restored the impaired proliferation under MTHFD2 inhibition to the level of vehicle-treated cells (Fig. 3B).

**Figure 3:**
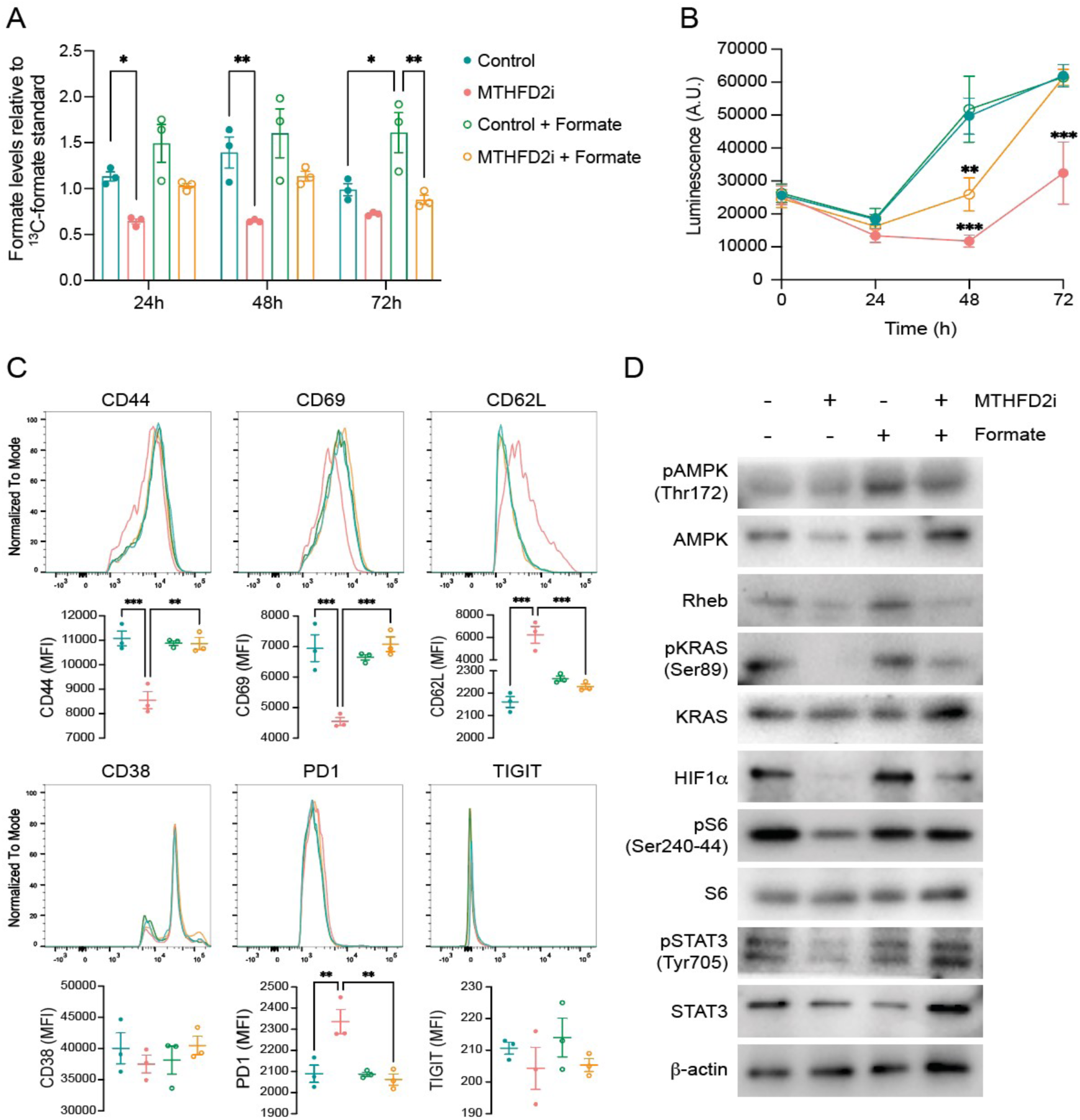
Effects of MTHFD2 inhibition on CD8⁺ T-cell response are mitigated by formate supplementation. (A) Intracellular formate levels relative to standard sodium ^13^C-formate, measured by GC-MS per 2×10^6^ cells, and (B) cell viability measured by CTG assay in activated CD8⁺ T cells treated with DMSO (control), 500 nM MTHFD2i, DMSO + 1 mM formate (control + formate), and 500 nM MTHFD2i + 1 mM formate (MTHFD2i + formate) for 24, 48, and 72 hours. (C) Flow cytometric analysis of the expression of activation (CD44, CD69), naïve (CD62L), and inhibitory (CD38, PD1, and TIGIT) markers (gated on non-stimulated cells), and (D) immunoblot of phospho-AMPK, AMPK, phospho-KRAS, KRAS, HIF1α, phospho-S6, S6, phospho-STAT3, STAT3, and b-actin (C) in activated CD8⁺ T cells treated with DMSO (control), 500 nM MTHFD2i, DMSO + 1 mM formate (control + formate), and 500 nM MTHFD2i + 1 mM formate (MTHFD2i + formate) for 48 hours. Activation was induced by anti-CD3 and -CD28 (aCD3/CD28) antibodies and IL-2. Data are shown as mean ± SEM. Statistical multiple comparison performed by one-way ANOVA (panel C) and two-way ANOVA (panel A and B): *p < 0.05, **p < 0.01, ***p < 0.001.

In accordance with our previous findings (Fig. 2), MTHFD2i reduced the activation of CD8^+^ T cells, as indicated by the upregulation of activation markers CD44 and CD69, along with the downregulation of L-selectin (CD62L). It also accelerated CD8^+^ T-cell exhaustion, as measured by the PD1 inhibitor marker (Fig. 3C). Consistently, formate alone did not enhance the T-cell response but reversed the effects of MTHFD2i (Fig. 3C). Additionally, formate partially addressed the deficiency of KRAS and the downstream mTOR targets S6, HIF1α, and STAT3 levels during MTHFD2i treatment (Fig. 3D), highlighting the essential role of formate in maintaining an adequate CD8^+^ T-cell response under impaired 1C mitochondrial metabolism.

### MTHFD2 inhibition lowers the TCF1^high^ CD8^+^ T-cell progenitor population and **b**-cell autoimmune reactivity

The glucose-6-phosphatase catalytic subunit-related protein (IGRP) is an islet-specific protein expressed in pancreatic β cells (Lieberman et al. 2003). The IGRP-reactive T-cell population constitutes the earliest islet infiltrates during T1D, and the response to IGRP may be one of the first events leading to β-cell destruction by diabetogenic CD8^+^ T cells (Shieh et al. 2005).

Recently, a study identified a specific population of IGRP-reactive CD8^+^ T-cell autoimmune progenitors that significantly induces T1D (Gearty et al. 2022). A critical feature of the IGRP CD8^+^ T cells is their unique phenotypic bifurcation into high and low expression levels of the transcription factor TCF1, which is critical for T-cell memory, longevity, and self-renewal (Zhang et al. 2021; Zhao, Shan, and Xue 2022). This bifurcation is highly relevant: as few as 100 TCF1^high^ cells are sufficient to drive T1D, while as many as 100,000 TCF1^low^ cells are not (Gearty et al. 2022).

Transcriptome analysis of TCF1^high^ versus TCF1^low^ cells showed that TCF1^high^ cells are enriched for pathways involved in DNA replication (Gearty et al. 2022). Given that MTHFD2 and formate play a crucial role in supplying nucleotides for DNA synthesis and repair via 1C metabolism, we investigated their potential impact on TCF1 expression in IGRP CD8^+^ T cells. We observed that MTHFD2i treatment decreases the percentage of TCF1^high^ cells in whole CD8^+^ T cells (Fig. 4A). When we sorted IGRP-specific CD8^+^ T cells using the tetramer NRP-V7, MTHFD2i consistently reduced the TCF1^high^ population and increased the TCF1^low^ population. Exogenous formate partially rescued this effect (Fig. 4B). Thus, MTHFD2 and formate may act as regulators of TCF1 in IGRP CD8^+^ T cells and, in doing so, regulate this key pathogenic CD8^+^ T-cell population in both mice and humans.

**Figure 4:**
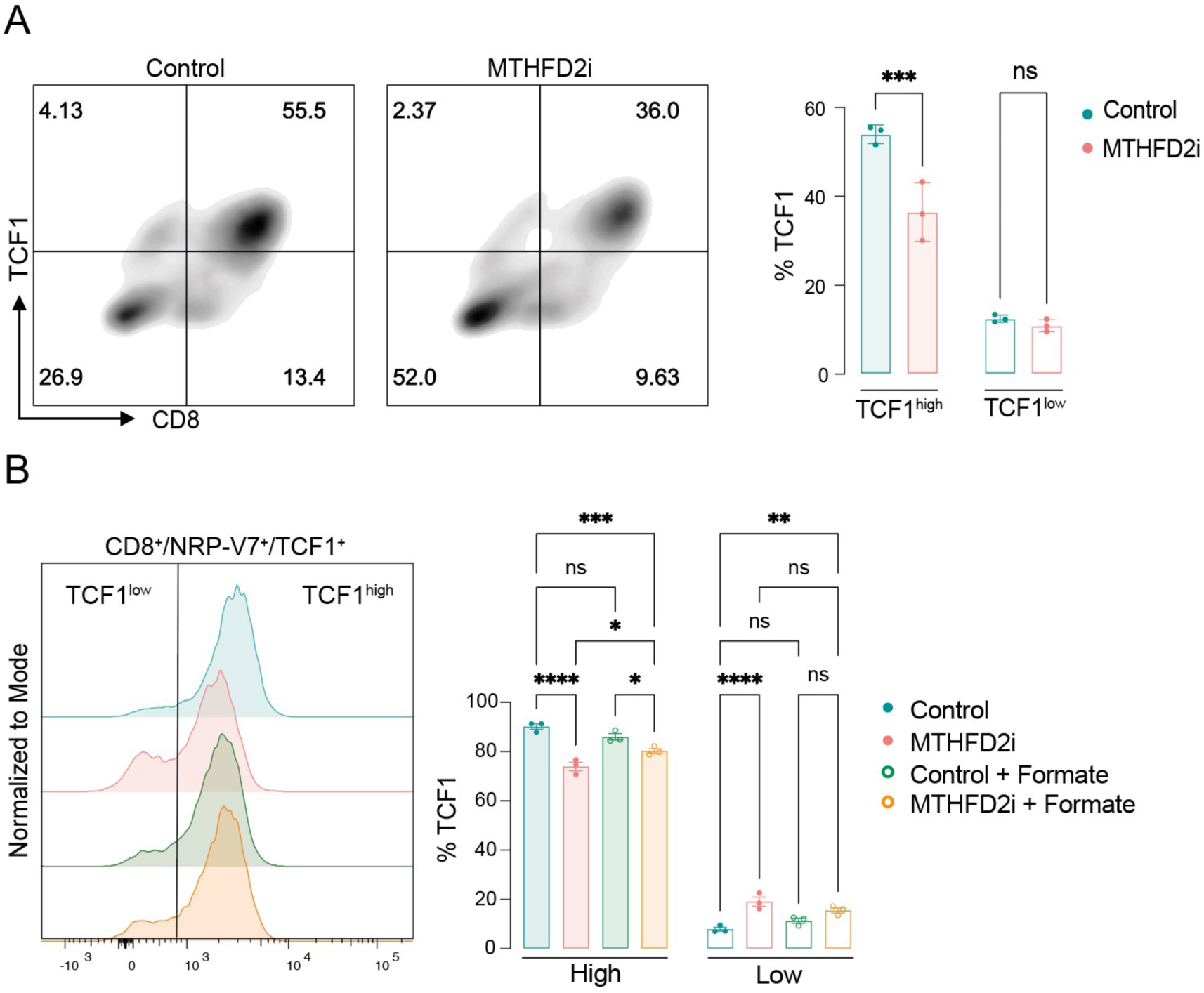
Inhibition of MTHFD2 shifts the proportion of TCF1^high^/TFC1^low^ stem-like autoimmune progenitor CD8⁺ T cells. (A) Flow cytometric analysis of the expression of the transcription factor T cell factor 1 (TCF1) in CD8⁺ T cells, activated by anti-CD3 and -CD28 (aCD3/CD28) antibodies and IL-2, and treated with DMSO (control) or 500 nM MTHFD2i for 48 hours. Top gates represent low (left) or high (right) TCF1 expression. (B) TCF1 expression in IGRP-specific CD8⁺ T cells under 48-hour activation with DMSO (control), 500 nM MTHFD2i, DMSO + 1 mM formate (control + formate), and 500 nM MTHFD2i + 1 mM formate (MTHFD2i + formate). Gates represent low (left) or high (right) TCF1 expression. Data are shown as mean ± SEM. Statistical comparisons between two groups in panel A were performed by unpaired t-test, and multiple comparisons in panel B were performed by one-way ANOVA: *p < 0.05, **p < 0.01, ***p < 0.001, ****p < 0.0001.

### MTHFD2 inhibition during the early insulitis stage delays T1D onset in NOD mice

We and others have demonstrated a clear effect of MTHFD2 inhibition on the modulation of immune-cell responses, including lymphocytes and macrophages. Thus, the therapeutic application of MTHFD2 inhibitors, extending beyond its anticancer roles to include protection against inflammation and autoimmunity, is gaining increasing prominence. MTHFD2 is consistently overexpressed in many human inflammatory and autoimmune diseases. Furthermore, it mitigates the severity of experimental autoimmune encephalomyelitis and protects against inflammatory bone loss in rheumatoid arthritis (Sugiura et al. 2022; Tang and Hou 2023; Yujing Li et al. 2025), suggesting a promising role of MTHFD2 in autoimmune conditions. However, its effect on autoimmune T1D has not been tested.

To investigate the role of MTHFD2 and formate in a T1D context, we used the NOD mouse model, in which, as in humans, unusual polymorphisms within an MHC class II molecule contribute the most to T1D risk (Driver, Chen, and Mathews 2012). The natural history of diabetes development in this model occurs in different stages: 1) infiltration of innate immune cells into the pancreas, including dendritic cells, macrophages, and neutrophils, without immune attack (within 3–4 weeks); 2) peri-insulitis, where innate immune cells attract adaptive CD4^+^ and CD8^+^ T cells into the pancreas, bordering the islet with minimal immune attack (within 8–10 weeks); 3) insulitis, characterized by complete invasion of CD4^+^ and CD8^+^ T cells into the islet and destruction of pancreatic β cells (within 12 weeks); and 4) onset of diabetes (within 16–35 weeks) (Fig. 5A) (Pearson, Wong, and Wen 2016; Thompson et al. 2019).

**Figure 5:**
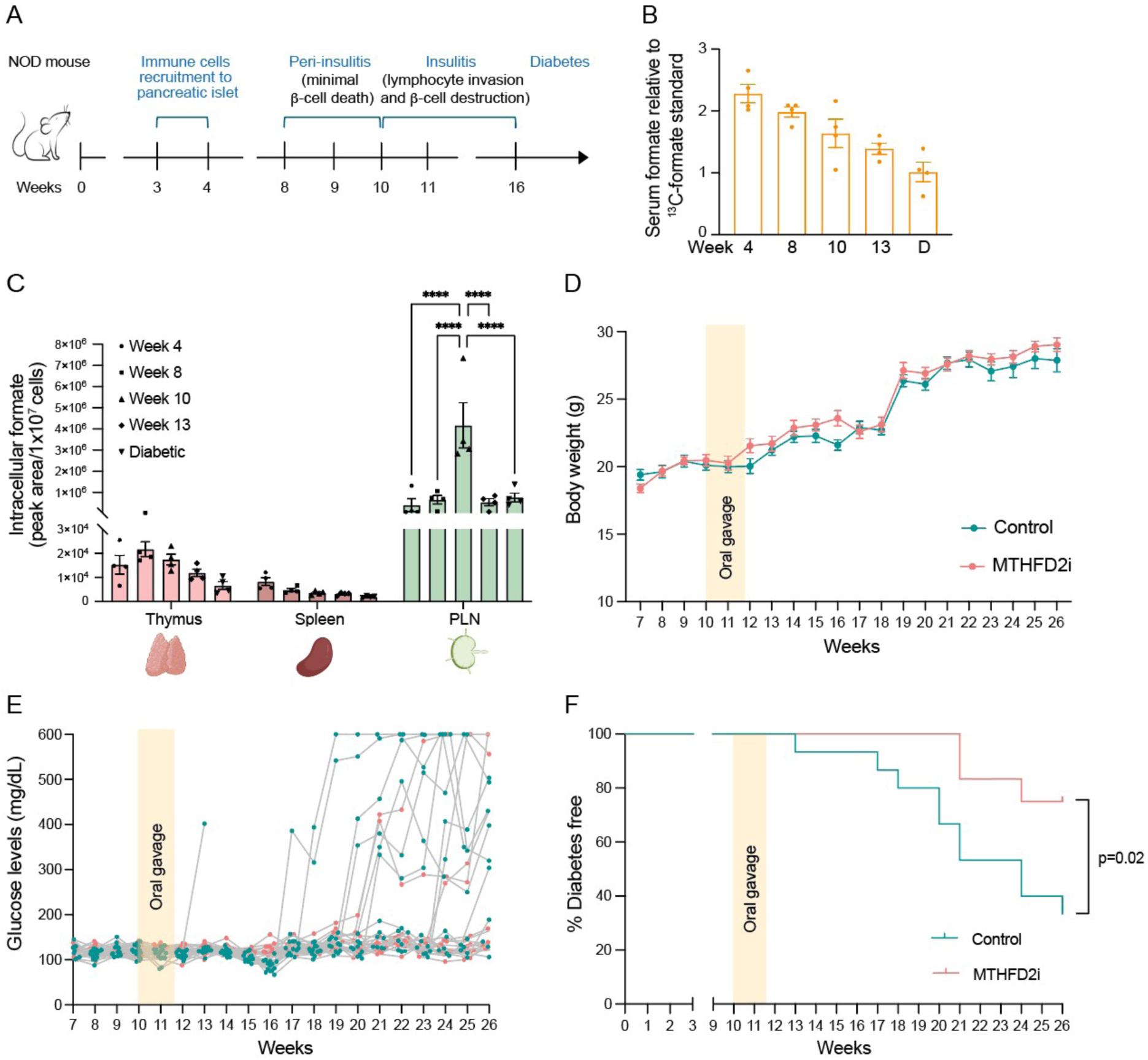
MTHFD2 inhibition during the early insulitis stage delays T1D onset in NOD mice. (A) Schematic of T1D pathogenesis stages in NOD mice. The development of diabetes in the NOD mouse model occurs in four stages: 1) lymphocyte recruitment to the pancreas without autoimmune attack (∼3–4 weeks), 2) peri-insulitis with minimal autoimmune attack (∼8–10 weeks), 3) insulitis with lymphocyte invasion and autoimmune pancreatic b-cell destruction (∼12– 16 weeks), and 4) diabetes onset (∼16–30 weeks). (B and C) Relative formate levels to standard sodium ^13^C-formate in 50 µL serum (B) and cellular thymus, spleen, and PLN (C), measured by GC-MS in euglycemic mice at 4, 8, 10, and 13 weeks and in diabetic NOD mice. Data are shown as mean ± SEM. (D–F) Body weight (D), blood glucose levels (E), and the percentage of nondiabetic mice (F) in a cohort of NOD mice treated twice daily with corn oil vehicle (n = 12) or 200 mg/kg MTHFD2 inhibitor (n = 15) orally for 10 days starting at week 10. Data are shown as mean ± SEM in panels B-D. Statistical analysis was performed by two-way ANOVA in panel C (****p < 0.0001) and log-rank test in panel F.

To understand the influence of formate on T1D development in the NOD model, we monitored serum levels to determine whether this metabolite could serve as a serological biomarker for disease prediction. Contrary to our expectations, we observed that serum levels of formate decreased with age and with the progression of T1D in NOD mice (Fig. 5B), suggesting that serum formate is unlikely to predict T1D.

CD4^+^ and CD8^+^ T cells are found not only in peripheral blood but also primarily in the thymus, spleen, and lymph nodes in mice (Hensel et al. 2019). Therefore, we examined the dynamics of formate in whole immune cells from these lymphoid tissues, which are essential for T-cell maturation and activation during T1D. Overall, we observed that intracellular formate levels are higher in the thymus and pancreatic lymph node (PLN) than in the spleen (Fig. 5C), correlating with tissues that have higher levels of mature CD4^+^ and CD8^+^ T cells and immature doublepositive CD4^+^/CD8^+^ T cells (Fig. S3). Additionally, we identified a formate peak in the thymus and PLN at 8 and 10 weeks of age, respectively, coinciding with the peri-insulitis and insulitis stages (Fig. 5C). This indicates a higher production of formate in immune cells during critical phases of T1D development.

Given these findings and the fact that MTHFD2i reduces formate levels *in vitro*, we assessed the impact of MTHFD2i on preventing or delaying T1D in the NOD mouse model. We treated the mice with the MTHFD2i starting at week 10, during the peak of formate in the PLN, for the maximum period of safety tested *in vivo* (10 days) (Kawai et al. 2019). Body weight remained similar between the control (corn oil) and MTHFD2i-treated groups throughout the 26-week monitoring period (Fig. 5D). Blood glucose levels were comparable prior to treatment (Fig. 5E). However, the control group maintained higher blood glucose levels (Fig. 5E), and diabetes onset occurred more rapidly than in the MTHFD2i group (Fig. 5F). At 15 weeks following MTHFD2i treatment, only 3 out of 12 mice developed diabetes, in contrast to 10 out of 15 mice in the control group. This finding suggests a potential new therapeutic role for MTHFD2 inhibition in delaying and preventing T1D.

## Discussion

Despite the central role of 1C metabolism in supplying 1C units that support multiple biological functions, and the extensive knowledge linking its disruption to various diseases such as cancer, its involvement in regulating the immune system is only now beginning to emerge (Kurniawan, Kobayashi, and Brenner 2021).

Recent studies have identified the mitochondrial 1C metabolism enzyme MTHFD2 as a control point for CD4^+^ T-cell responses (Sugiura et al. 2022). Due to its effects in autoimmune encephalomyelitis and rheumatoid arthritis, MTHFD2 has become a highly attractive therapeutic target for autoimmune diseases (Sugiura et al. 2022; Yujing Li et al. 2025). However, the role of MTHFD2 and its product, the metabolite formate, in CD8^+^ T cells within the context of T1D remains unclear. Our observations suggest that inhibiting MTHFD2 and formate production regulates not only the expansion but also the activation and effector functions of CD8^+^ T cells. Furthermore, targeting MTHFD2 in T cells delays and reduces T1D onset in NOD mice, highlighting this enzyme as a promising new target for T1D treatment.

Consistent with other reports (Ron-Harel et al. 2016; Ma et al. 2017), we found that 1C metabolism enzymes, including MTHFD2, are upregulated during the CD8^+^ T-cell response. Disrupting this pathway, either by inhibition of MTHFD2, dual inhibition of the serine hydroxymethyltransferases SHMT1 and SHMT2, or serine and glycine starvation, significantly reduces CD4^+^ and CD8^+^ T-cell proliferation (Ma et al. 2017; Ron-Harel et al. 2016). Furthermore, we found that MTHFD2 inhibition not only impairs CD8^+^ T-cell activation but may also affect two key features of CD8^+^ T cells in T1D: exhaustion and self-renewal.

In the context of cancer or chronic viral infection, CD8^+^ T cells enter a hyporesponsive state referred to as T-cell exhaustion or dysfunction after persistent antigen stimulation (Tonnerre et al. 2021; Gearty et al. 2022). However, in autoimmunity, despite chronic self-antigen exposure, CD8^+^ T cells do not become dysfunctional; instead, they maintain effector functions and progressively destroy the tissue (Gearty et al. 2022). Thus, in T1D, CD8^+^ T-cell exhaustion is linked to positive therapeutic outcomes (Wiedeman et al. 2020; Linsley and Long 2019). In fact, clinical studies have shown that the anti-CD3 monoclonal antibody teplizumab exerts beneficial effects in T1D by promoting CD8^+^ T-cell exhaustion (Alice Long et al. 2016; Herold et al. 2024). Exhausted CD8^+^ T cells typically express inhibitory receptors like programmed cell death 1 (PD1) (Tonnerre et al. 2021), and we observed that inhibiting MTHFD2 increases PD1 expression. Although CD8^+^ T-cell exhaustion due to chronic stimulation can be mitigated, stem-like, antigen-specific T cells can offset the loss of CD8^+^ T cells and serve as an essential long-term reservoir for autoimmune CD8^+^ T-cell responses (Aljobaily et al. 2024). Furthermore, stem-like CD8^+^ T cells, defined by expression of the transcription factor TCF1, drive T1D (Gearty et al. 2022). Our findings show that MTHFD2 inhibition not only promotes CD8^+^ T-cell exhaustion but also decreases TCF1 expression in the IGRP-specific CD8^+^ T-cell population, indicating that targeting MTHFD2 could be a potential alternative strategy for autoimmunity with dual functions for managing CD8^+^ T-cell exhaustion and self-renewal.

Because TCF1 is a known target of canonical Wnt signaling (Gearty et al. 2022), our results indicate a link between MTHFD2 and the Wnt pathway. Although various transcriptional signals, including mTORC1, MYC, or Wnt/β-catenin, can regulate MTHFD2 expression (Ben-Sahra et al. 2016; Zhu and Leung 2020; Pardo-Lorente and Sdelci 2024), whether MTHFD2 directly affects these factors and the mechanisms behind these connections remain unclear.

Many of the effects observed with MTHFD2 inhibition in CD8^+^ T cells, such as suppression of cell expansion, likely stem from its canonical enzymatic functions, which supply intermediates necessary for *de novo* purine synthesis and formylation reactions (Pardo-Lorente and Sdelci 2024). Downstream, MTHFD2 deficiency can promote AICAR accumulation, guanine depletion, and subsequent inhibition of the mTORC1-activating GTPase Rheb, thereby suppressing mTORC1 (Kim et al. 2016; Emmanuel et al. 2017; Sugiura et al. 2022).

Although inhibiting MTHFD2 reduced mTORC1 downstream targets (i.e., S6, HIF1α, and STAT3), these effects were not associated with changes in AICAR or Rheb levels. Instead, MTHFD2 inhibitor treatment appeared to modulate mTORC1 targets by suppressing KRAS. A previous report indicated that depletion of KRAS decreases MTHFD2 expression in colorectal cancer cells and that KRAS transcriptionally upregulates MTHFD2 (Ju et al. 2019). Our results suggest a new function of MTHFD2 in suppressing KRAS in CD8^+^ T cells. However, further investigation is needed to determine whether MTHFD2 directly interacts with or regulates KRAS.

Together with MTHFD2, formate, the product of the mitochondrial branch of 1C metabolism, plays a crucial role in determining the downstream functions of 1C metabolism by serving as a 1C donor feeding into cytosolic and nuclear nucleotide synthesis and methylation reactions (Lamarre et al. 2013; Green et al. 2023). Despite these essential functions, few studies have addressed the direct impact of formate on T cells.

We revealed for the first time the dynamics of formate levels during CD4^+^ and CD8^+^ T-cell activation and identified a peak in formate production 48 hours post-activation *in vitro*. Additionally, we demonstrated that the inhibition of MTHFD2 reduces intracellular formate levels. Exogenous formate can rescue MTHFD2, SHMT1, and SHMT2 inhibition as well as serine starvation effects on T-cell proliferation *in vitro* (Ron-Harel et al. 2016; Ma et al. 2017; Sugiura et al. 2022; Rowe et al. 2023) and improve the effectiveness of anti–PD1 therapies in CD8^+^ T-cell cancer immunotherapy *in vitro* and *in vivo* (Rowe et al. 2023). Our results show that although formate supplementation restores the CD8^+^ T-cell proliferation, activation, and function impaired by MTHFD2 inhibition, it does not further enhance the CD8^+^ T-cell responses when administered alone. However, exogenous formate prevented the loss of the stem-like CD8^+^ T-cell phenotype driven by MTHFD2 inhibition *in vitro*, suggesting a new potential role for this metabolite. Given that CD8^+^ T cells are the most abundant lymphocytes in islets during the insulitis stage and are primarily responsible for β-cell destruction during T1D (Wiedeman et al. 2020; Gearty et al. 2022), we extended our *in vitro* findings to the *in vivo* NOD mouse model.

Considering that the insufficient availability of biomarkers represents a significant gap in understanding the cause and progression of T1D (Nakayasu et al. 2023), we hypothesized that formate could represent a promising new biomarker for the disease. Although we did not observe elevated serum formate levels during the development of T1D, measurement of formate in lymphoid organs revealed higher concentrations in the PLN tissue, where autoreactive T cells become activated and from which effector T cells migrate to the pancreas (Marrero et al. 2012). We also observed a peak in formate at the onset of insulitis, showing for the first time a pivotal role for formate in the development of T1D. Furthermore, inhibiting MTHFD2 and consequently lowering formate levels delayed disease onset and reduced the incidence of T1D in the NOD mouse model.

In sum, our work emphasizes the importance of mitochondrial 1C metabolism in the CD8^+^ T-cell response and provides evidence supporting this pathway as a therapeutic target for autoimmune conditions such as T1D. Clinically used, FDA-approved drugs that target 1C metabolism, such as the anti-folate MTX, have not improved outcomes in T1D and have been associated with an earlier increase in insulin requirements (Buckingham and Sandborg 2000). Our findings suggest that inhibiting MTHFD2 and its product, formate, could establish a new paradigm for T1D clinical management and pave the way for innovative approaches to treating T1D. Determining the direct effect of MTHFD2 inhibitors on CD8^+^ T cells *in vivo* and optimizing the timing and duration of treatment remain important challenges to address.

## Materials and Methods

### Mice

The use of animals was approved by the Harvard Medical School (HMS) Institutional Animal Care and Use Committee (IACUC, no. IS00003204). Wild-type C57BL/6J and NOD/ShiLtJ mice were purchased from The Jackson Laboratory and housed at the Harvard Center for Comparative Medicine, on a regular light cycle (12-h dark/light), 22–24°C, 40–60% humidity, with free access to water and a standard laboratory chow diet (PicoLab Mouse Diet 5058, LabDiet). Female C57BL/6J mice between 8 and 10 weeks old were used for *in vitro* studies. Female euglycemic (between 3–13 weeks old) and diabetic (older than 13 weeks old) NOD/ShiLtJ mice were used for *in vivo* studies.

### Key antibodies and reagents Immunoblotting

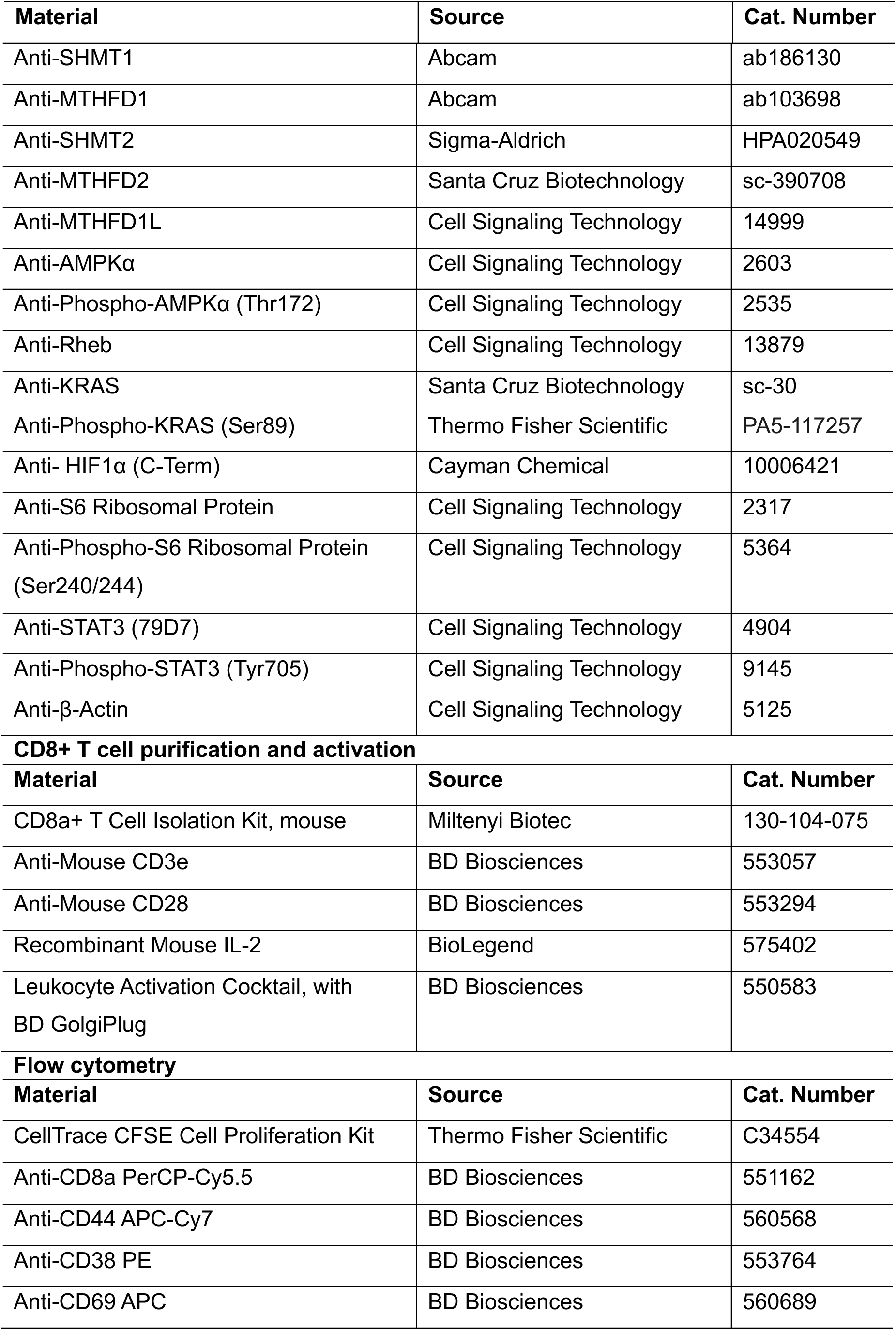
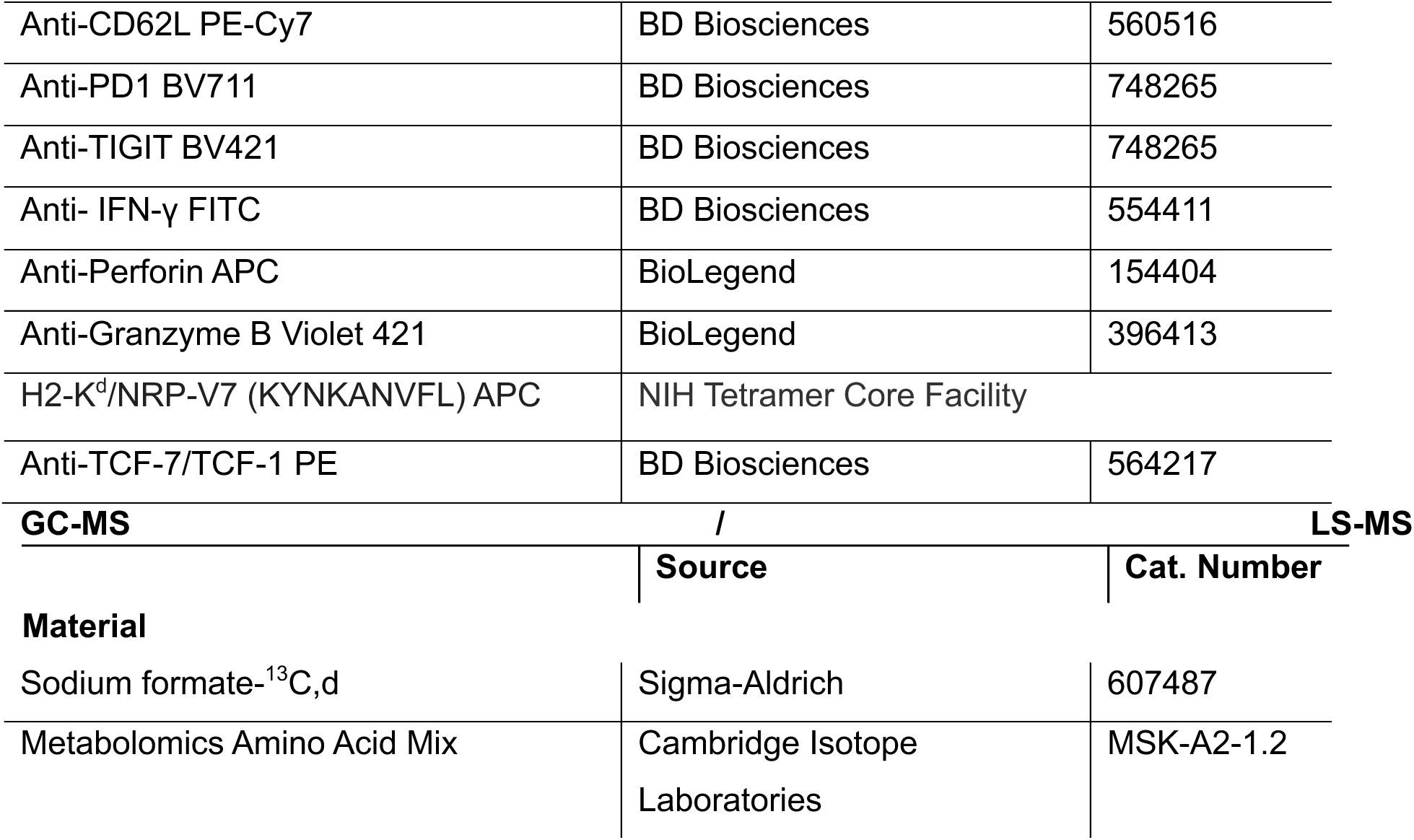

### Mouse immune cell isolation

Primary mouse T-cell isolation was conducted as previously described (Grosjean et al. 2021). Briefly, female mice were anesthetized with isoflurane, euthanized by cervical dislocation, and the abdominal cavity was opened. The spleen was removed, placed in cold T-cell buffer (PBS, 0.5% BSA, 2 mM EDTA), minced with the piston of a syringe, and passed through a 70-μm cell strainer. Red blood cells were lysed with RBC lysis buffer (Thermo Fisher Scientific, 00-4333-57) for 8 min at 37°C. The reaction was stopped by doubling the volume with T-cell buffer. The final pellet was resuspended in 60 μL T-cell buffer for CD8⁺ T-cell purification.

### Mouse CD8**⁺** T-cell purification, activation, and culture conditions

Primary mouse CD8⁺ T cells were purified from the spleen using a negative isolation kit following the manufacturer’s instructions. After purification, cells were counted and cultured at 37°C with 5% CO₂ in RPMI media (Sigma-Aldrich, 1640), supplemented with 10% dialyzed FBS, 10 mM HEPES, and 50 μM 2-mercaptoethanol. Activation was induced with soluble anti-CD3 (2 μg/mL), anti-CD28 (4 μg/mL), and IL-2 (10 ng/mL). Cells were activated in the presence of either DMSO control or DS18561882 MTHFD2 inhibitor (MedChemExpress, HY-130251) at a 500 μM dose. Formate rescue experiments were conducted with 1 mM sodium formate. Cells were seeded at 1.5×10⁵ cells/well for a 96-well plate, 1×10⁶ cells/well for a 24-well plate, or 2×10⁶ cells/well for a 6-well plate for 0–72 hours.

### Immunoblotting

Activated primary mouse CD8⁺ T cells cultured under specific experimental conditions (±MTHFD2i / ±formate) for 0–72 hours were harvested, washed with PBS, and lysed using RIPA Lysis and Extraction Buffer (Thermo Fisher Scientific, 89901) and a Protease and Phosphatase Inhibitor Cocktail (Thermo Fisher Scientific, 78446). Protein concentration was quantified using a BCA Protein Assay (Thermo Fisher Scientific, 23227). Between 10–20 μg of protein was loaded for polyacrylamide gel electrophoresis with Tris-Glycine Mini precast gels (4–20%) (Thermo Fisher Scientific, XP04205BOX). The resolved gels were transferred to PVDF membranes and blocked with 5% non-fat dry milk in Tris-buffered saline containing 0.05% Tween-20 (TBS-T) before incubation with the following primary antibodies: β-Actin, SHMT1, MTHFD1, SHMT2, MTHFD2, MTHFD1L, AMPKα, pAMPKα (Thr172), Rheb, KRAS, pKRAS (Ser89), HIF1α, S6, pS6, STAT3, and pSTAT3 (Tyr705), for 1–2 hours at room temperature or overnight at 4°C. All antibodies were used at a dilution of 1:1000. Membranes were then incubated with HRPconjugated secondary antibody (1:3000) for 1–2 hours at room temperature, visualized using Pico chemiluminescent substrate (Thermo Fisher Scientific, 34580), and imaged with the ChemiDoc MP Imaging System, Version 6.0 (Bio-Rad, 2011).

### Formate extraction and derivatization for GC-MS analysis

#### *Analysis of formate in cultured primary mouse CD8*⁺ *T cells*

Naïve or activated primary mouse CD8⁺ T cells were cultured under specific experimental conditions (±MTHFD2i / ±formate) for 0–72 hours. Metabolite extraction was conducted as previously described (Meiser et al., 2016). Briefly, 2×10⁶ cells were counted, collected, and centrifuged at 350g for 5 min. The cell pellet and media were separated and processed. The cell pellet was washed with cold PBS and incubated with cold extraction solution (30% methanol / 20% acetonitrile / 50% water, chromatography grade, 400 μL per 1×10⁶ cells) on a rocking shaker at 4°C for 10 min. For the media, 10 μL of sample was diluted in 990 μL of cold extraction solution. For both cases, samples were vortexed for 1 min and centrifuged at 15,000g for 5 min at 4°C. The supernatant was transferred into GC-MS glass vials with inserts and stored at −80°C until formate derivatization and analysis.

Formate was derivatized to pentafluorobenzyl formate for detection by GC-MS following the protocol previously described (Lamarre et al. 2014). 50 μL of sample and 20 μL of standard sodium ¹³C-formate, prepared in 0.5 M phosphate buffer (pH 8), were mixed with 130 μL of 2,3,4,5,6-pentafluorobenzyl bromide (PFBBr) in acetone (alkaline conditions). Samples were vigorously vortexed for at least 1 min and incubated at 60°C for 15 min. Extraction was performed using 330 μL of n-hexane, and 200 μL of the organic phase was transferred into GC-MS glass vials with inserts and analyzed by GC-MS the same day.

Derivatized formate samples were analyzed using a DB-FastFAME column (Agilent Technologies, G3903-63011) in an Agilent 7890A gas chromatograph coupled with an Agilent 5975C mass spectrometer. The *m/z* was monitored at 226 and 227, representing the ¹²C-formate and ¹³C-formate derivatives, respectively.

#### Analysis of formate in serum and immune cells from lymphoid organs

Blood, thymus, spleen, and PLN were collected from euglycemic (4-, 8-, 10-, and 13-week-old) and diabetic NOD mice. Blood was centrifuged for 15 min at 3,000 rpm at 4°C, and the serum (supernatant) was collected and stored at −20°C until analysis. For the thymus, spleen, and PLN, immune cell isolation and metabolite extraction were performed as described above. Cells were counted and processed for metabolite extraction. A volume of 50 μL of the direct serum or cell metabolite extraction sample was derivatized and analyzed by GC-MS as indicated previously.

Relative quantitation of formate was conducted using El-MAVEN (v.0.12.0) with a 20 ppm mass tolerance. Formate measurements were normalized to the labeled ¹³C-formate standard and to cell number.

### Cell viability and proliferation

Cell counting and viability were assessed by mixing a final volume of 10 μL of cell suspension with 0.4% Trypan Blue Solution (Thermo Fisher Scientific, 15250061) and loading into a slide counter for automated counting (Bio-Rad, TC20).

Cell proliferation was assessed by quantifying ATP content in viable cells via luminescence using the CellTiter-Glo (CTG) Assay (Promega, G7570). In brief, 1.5×10⁵ cells/well, in a final volume of 50 μL, were seeded, activated, and cultured under the specific experimental conditions (±MTHFD2i / ±formate) in 96-well plates. After 0, 24, 48, and 72 hours, 50 μL of CTG reagent was added, and luminescence was measured.

For the CFSE proliferation assay, a CellTrace Kit (Thermo Fisher Scientific, C34554) was used according to the manufacturer’s instructions. CD8⁺ T cells were purified and incubated in 1 μM CellTrace CFSE staining solution for 10 min at 37°C. The reaction was halted by quintupling the volume with medium. Cells were spun down and cultured under the specific experimental conditions (±MTHFD2i / ±formate). Dye dilution was measured by flow cytometry 48 hours later.

### Surface and intracellular staining for flow cytometric analysis

Activated primary mouse CD8⁺ T cells were cultured under specific experimental conditions (±MTHFD2i / ±formate) for 48 hours. Surface marker staining was performed by collecting, washing, and incubating cells with antibodies against CD8, CD44, CD69, CD62L, CD38, PD1, and TIGIT for 30 min at 4°C, diluted 1:500 each in T-cell buffer (PBS, 0.5% BSA, 2 mM EDTA). The cells were then spun down, washed, and resuspended in 400 μL T-cell buffer for flow cytometry analysis.

Before intracellular cytokine and nuclear factor staining, cells were re-stimulated with the phorbol ester PMA and ionomycin for 4 hours at 37°C in the presence of GolgiPlug (BD Biosciences, 550583). Afterward, cells were collected, washed, stained first with surface markers (e.g., CD8), and then fixed and permeabilized using the eBioscience Foxp3/Transcription Factor Staining Buffer Set (Invitrogen, 00-5523-00), following the manufacturer’s protocol. Intracellular staining was performed by incubating cells with antibodies against IFNγ, Perforin, Granzyme B, NRP-V7, and TCF1 (1:100) for 20–30 min at room temperature. The cells were then spun down, washed, and resuspended in 400 μL T-cell buffer for flow cytometry analysis.

Flow cytometric analysis was performed using an LSRFortessa and BD FACSDiva 8.0.1 software at the Harvard School of Public Health, Department of Molecular Metabolism core instrumentation. Flow data were analyzed with FlowJo v.10 software and gated based on nonstimulated (naive) CD8^+^ T cells (Fig. S4).

### Mass spectrometry for nucleotide intermediate analysis

Nucleotide intermediate levels were measured by LC-MS as previously described (Kory et al. 2018; Diehl et al. 2022). Briefly, 1×10⁶ activated primary mouse CD8⁺ T cells were cultured under the specific experimental conditions (±MTHFD2i / ±formate) for 48 hours and re-stimulated with the phorbol ester PMA and ionomycin for 4 hours at 37°C in the presence of GolgiPlug (BD Biosciences, 550583). Polar metabolites were extracted in 500 μL of ice-cold 80% methanol in water with 250 nM ¹³C/¹⁵N-labeled amino acid standards. Samples were vortexed for 10 min at 4°C and centrifuged at 17,000 × g. The supernatant was dried by vacuum centrifugation at 4°C, and the pellet was resuspended in 25 μL of a 50/50 acetonitrile/water mixture. Metabolites were then analyzed by LC-MS as previously described (Diehl et al. 2022).

### MTHFD2 inhibitor treatment in NOD model

Ten-week-old NOD mice were separated into two groups: control (n = 15) and MTHFD2i (n = 12). Both groups received oral gavage twice daily (8 a.m./8 p.m.) for 10 days. The control group was treated with corn oil, while the MTHFD2i group received the DS18561882 inhibitor at a dose of 200 mg/kg.

### Body weight, glucose measurement, and diabetes assessment

Blood glucose levels of mice (using a Roche Accu-Chek glucose meter) and body weight were monitored at least weekly in the morning, without fasting. Mice were considered diabetic after three consecutive days of blood glucose measurement >250 mg/dL.

### Pancreas histology and insulitis scoring

Control and MTHFD2i-treated diabetic NOD mice were sacrificed, and the pancreas was collected and fixed in 4% paraformaldehyde (Santa Cruz Biotechnology, sc-281692). Fixed pancreatic tissues were embedded in paraffin, sectioned at 5 μm thickness, and stained with hematoxylin and eosin (H&E) for histological examination of immune infiltration in pancreatic islets by the Rodent Histopathology Core at the Dana-Farber/Harvard Cancer Center facility based at HMS. Insulitis was scored as follows: 0, no infiltrate; 1, peri-islet infiltrate (25% islet destruction); 2, intraislet infiltrate (50% islet destruction); 3, intra-islet infiltrate (50–75% islet destruction); 4, complete islet destruction (Gearty et al. 2022; Kong et al. 2021).

### Statistical analysis

Statistical analyses were conducted using Prism software (Version 10.2.3) for unpaired two-tailed *t* test, one- or two-way ANOVA, and log-rank test, as appropriate. A p-value of <0.05 was considered statistically significant. All experiments were performed with at least three biological replicates.

## Author Contributions

G.R.H. and N.K. conceptualized the study. G.R.H., M.B., B.K., and S.B. performed experiments and analyzed data. S.B. and M.G.V.H. were instrumental in establishing and carrying out formate measurements. G.R.H wrote the manuscript and N.K. edited it with input from all authors.

## Acknowledgements

We thank Dr. Smita Gopinath and all members of the Kory lab for thoughtful discussions and feedback, and Dr. Julie Gosse for editing the manuscript. M.G.V.H. acknowledges support from R35CA242379 and P30CA014051. This work was supported by the Secretaría de Educación, Ciencia, Tecnología e Innovación de la Ciudad de México (SECTEI/130/2023) fellowship to G.R.H., and R35/MIRA (GM151097), a Damon-Runyon Rachleff Innovation Award (73-22), a JDRF Innovative award (1-INO-2023-1342-A-N), to N.K., and a gift by Richard Ranti.

## Supplementary Figures

**Figure S1:**
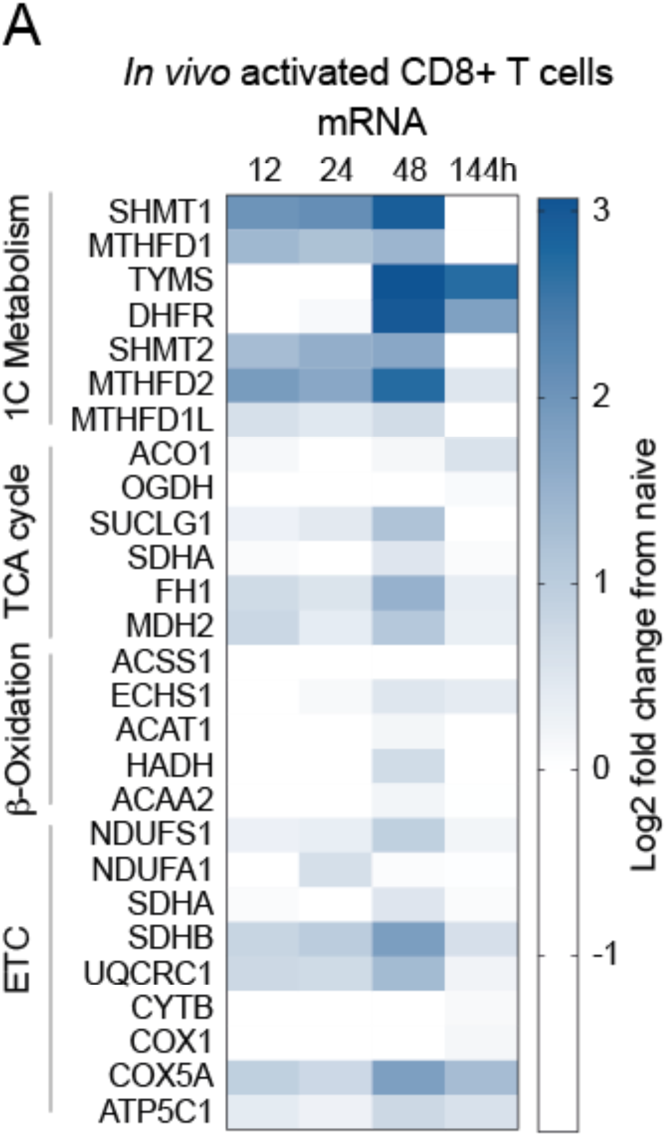
Transcriptome analysis of mitochondrial metabolic pathways in CD8+ T cells. (A) Heatmap of RNA sequencing displaying the fold change of individual key genes within the main mitochondrial metabolic pathways during the *in vivo* (Lm-OVA infection) compared to naïve cells.

**Figure S2:**
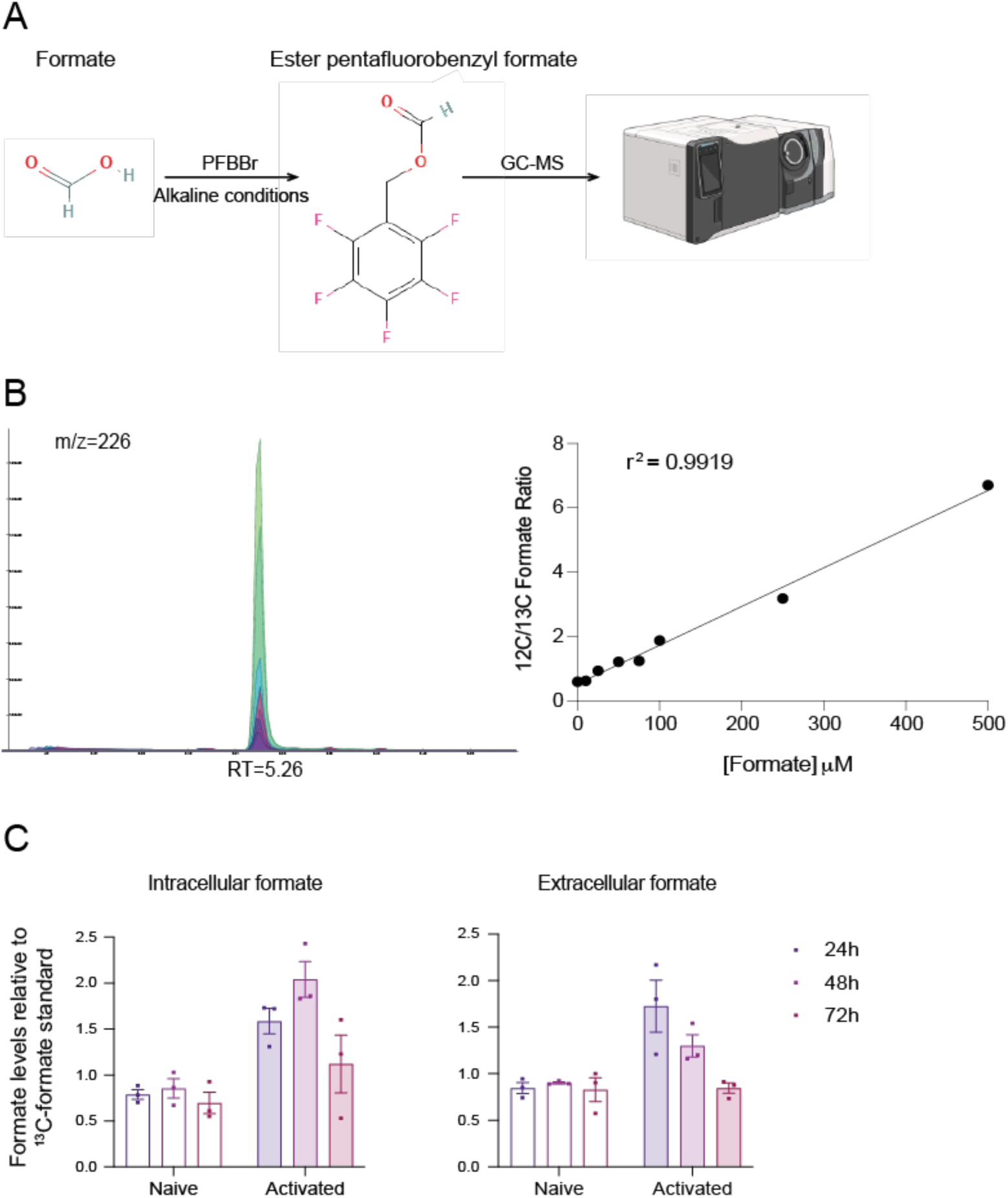
Quantification of formate by GC-MS. (A) Schematic of formate derivatization with pentafluorobenzyl bromide (PFBBr), forming the PFB formate ester for detection and quantification by GC-MS. (B) Representative chromatogram and standard curve from 0, 10, 25, 50, 75, 100, 250, and 500 mM of sodium formate measured by GC-MS using ^13^C-formate as an internal standard and monitoring *m/z* at 226 for ^12^C-formate and 227 for ^13^C-formate. (C) Intracellular and extracellular formate levels per 2×10^6^ cells relative to standard sodium ^13^Cformate measured by GC-MS in naïve and activated CD4⁺ T cells. Data are shown as mean ± SEM.

**Figure S3:**
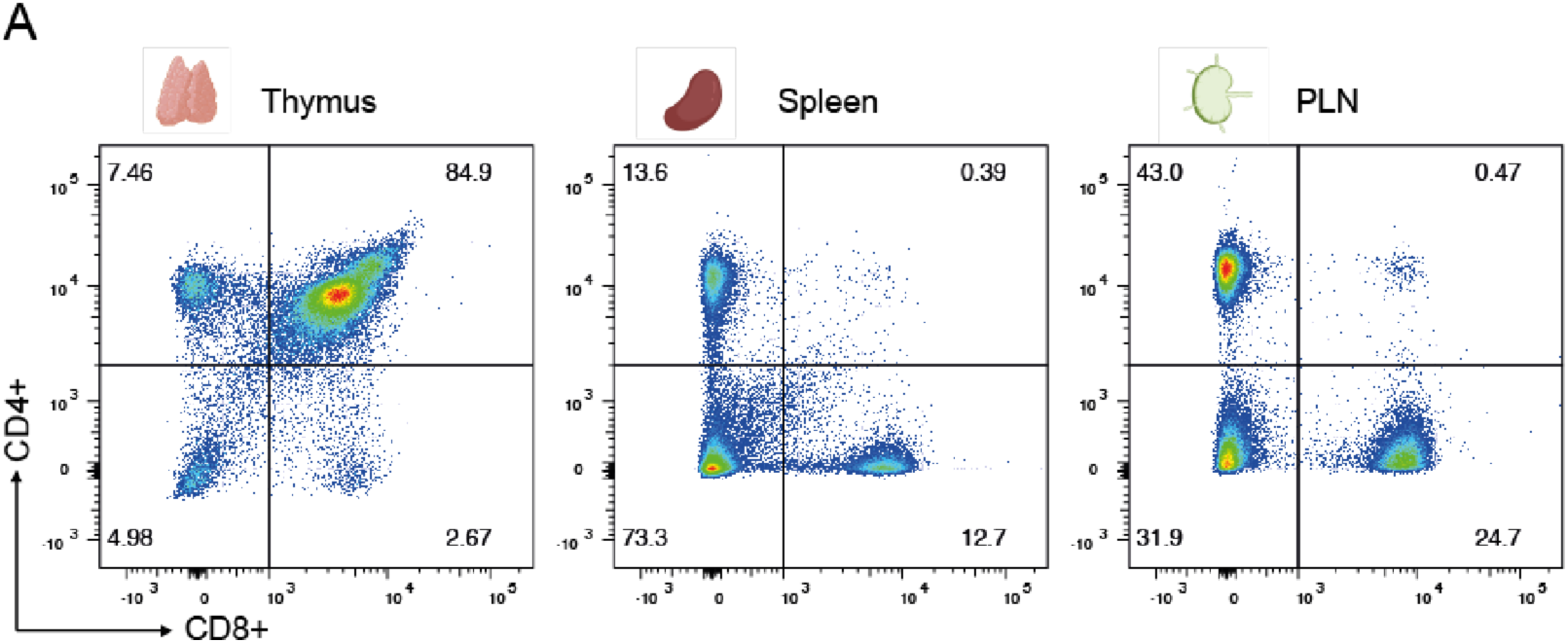
Proportion of CD4^+^ and CD8^+^ T cells in mouse lymphoid organs. (A) CD4⁺ and CD8⁺ expression in immune cells from the mouse thymus, spleen, and PLN, measured by flow cytometry in euglycemic 10-week-old NOD mice.

## References

Agarwal, Stuti, Catherine M. Bell, Scott B. Rothbart, and Richard G. Moran. 2015. “AMPActivated Protein Kinase (AMPK) Control of MTORC1 Is P53- and TSC2-Independent in Pemetrexed-Treated Carcinoma Cells.” Journal of Biological Chemistry 290 (46): 27476–86. 10.1074/jbc.M115.665133.

Alice Long, S, Jerill Thorpe, Hannah A DeBerg, Vivian Gersuk, James A Eddy, Kristina M Harris, Mario Ehlers, Kevan C Herold, Gerald T Nepom, and Peter S Linsley. 2016. “Partial Exhaustion of CD8 T Cells and Clinical Response to Teplizumab in New-Onset Type 1 Diabetes.” https://www.science.org.

Aljobaily, Nouf, Denise Allard, Bryant Perkins, Arielle Raugh, Tessa Galland, Yi Jing, W. Zac Stephens, Matthew L. Bettini, J. Scott Hale, and Maria Bettini. 2024. “Autoimmune CD4+ T Cells Fine-Tune TCF1 Expression to Maintain Function and Survive Persistent Antigen Exposure during Diabetes.” *Immunity*, November. 10.1016/j.immuni.2024.09.016.

Ben-Sahra, Issam, Gerta Hoxhaj, Stéphane J.H. Ricoult, John M. Asara, and Brendan D. Manning. 2016. “MTORC1 Induces Purine Synthesis through Control of the Mitochondrial Tetrahydrofolate Cycle.” Science 351 (6274): 728–33. 10.1126/science.aad0489.

Best J. Adam, David A. Blair, Jamie Knell, Edward Yang, Viveka Mayya, Andrew Doedens, Michael L. Dustin, et al. 2013. “Transcriptional Insights into the CD8 + T Cell Response to Infection and Memory T Cell Formation.” Nature Immunology 14 (4): 404–12. 10.1038/ni.2536.

Buckingham, Bruce A., and Christy I. Sandborg. 2000. “A Randomized Trial of Methotrexate in Newly Diagnosed Patients with Type 1 Diabetes Mellitus.” Clinical Immunology 96 (2): 86–90. 10.1006/clim.2000.4882.

Chapman, Nicole M., Mark R. Boothby, and Hongbo Chi. 2020. “Metabolic Coordination of T Cell Quiescence and Activation.” Nature Reviews Immunology. Nature Research. 10.1038/s41577-019-0203-y.

Debuysschere, Cyril, Magloire Pandoua Nekoua, Enagnon Kazali Alidjinou, and Didier Hober. 2024. “The Relationship between SARS-CoV-2 Infection and Type 1 Diabetes Mellitus.” Nature Reviews Endocrinology. Nature Research. 10.1038/s41574-02401004-9.

Diehl, Frances F., Teemu P. Miettinen, Ryan Elbashir, Christopher S. Nabel, Alicia M. Darnell, Brian T. Do, Scott R. Manalis, Caroline A. Lewis, and Matthew G. Vander Heiden. 2022. “Nucleotide Imbalance Decouples Cell Growth from Cell Proliferation.” Nature Cell Biology 24 (8): 1252–64. 10.1038/s41556-022-00965-1.

Driver, John P., Yi Guang Chen, and Clayton E. Mathews. 2012. “Comparative Genetics: Synergizing Human and NOD Mouse Studies for Identifying Genetic Causation of Type 1 Diabetes.” Review of Diabetic Studies. Society for Biomedical Diabetes Research. 10.1900/RDS.2012.9.169.

Ducker, Gregory S., Li Chen, Raphael J. Morscher, Jonathan M. Ghergurovich, Mark Esposito, Xin Teng, Yibin Kang, and Joshua D. Rabinowitz. 2016. “Reversal of Cytosolic One-Carbon Flux Compensates for Loss of the Mitochondrial Folate Pathway.” Cell Metabolism 23 (6): 1140–53. 10.1016/j.cmet.2016.04.016.

Ducker, Gregory S., and Joshua D. Rabinowitz. 2017. “One-Carbon Metabolism in Health and Disease.” Cell Metabolism. Cell Press. 10.1016/j.cmet.2016.08.009.

Dwyer, Alexander J., Jacob M. Ritz, Jason S. Mitchell, Tijana Martinov, Mohannad Alkhatib, Nubia Silva, Christopher G. Tucker, and Brian T. Fife. 2021. “Enhanced CD4+ and CD8+ T Cell Infiltrate within Convex Hull Defined Pancreatic Islet Borders as Autoimmune Diabetes Progresses.” Scientific Reports 11 (1). 10.1038/s41598-021-96327-2.

Emmanuel, Natasha, Shoba Ragunathan, Qin Shan, Fang Wang, Andreas Giannakou, Nanni Huser, Guixian Jin, Jeremy Myers, Robert T. Abraham, and Keziban Unsal-Kacmaz. 2017. “Purine Nucleotide Availability Regulates MTORC1 Activity through the Rheb GTPase.” Cell Reports 19 (13): 2665–80. 10.1016/j.celrep.2017.05.043.

Gassaway, Brandon M., Edward L. Huttlin, Emily M. Huntsman, Tomer M. Yaron-Barir, Jared L. Johnson, Kiran Kurmi, Lewis C. Cantley, et al. 2024. “Profiling Proteins and Phosphorylation Sites During T Cell Activation Using an Integrated Thermal Shift Assay.” Molecular & Cellular Proteomics: MCP 23 (7): 100801. 10.1016/j.mcpro.2024.100801.

Gearty, Sofia V., Friederike Dündar, Paul Zumbo, Gabriel Espinosa-Carrasco, Mojdeh Shakiba, Francisco J. Sanchez-Rivera, Nicholas D. Socci, et al. 2022. “An Autoimmune Stem-like CD8 T Cell Population Drives Type 1 Diabetes.” Nature 602 (7895): 156–61. 10.1038/s41586-021-04248-x.

Green, Alanna C., Petra Marttila, Nicole Kiweler, Christina Chalkiadaki, Elisée Wiita, Victoria Cookson, Antoine Lesur, et al. 2023. “Formate Overflow Drives Toxic Folate Trapping in MTHFD1 Inhibited Cancer Cells.” Nature Metabolism 5 (4): 642–59. 10.1038/s42255-023-00771-5.

Grosjean, Clémence, Julie Quessada, Mathis Nozais, Marie Loosveld, Dominique Payet-Bornet, and Cyrille Mionnet. 2021. “Isolation and Enrichment of Mouse Splenic T Cells for Ex Vivo and in Vivo T Cell Receptor Stimulation Assays.” STAR Protocols 2 (4). 10.1016/j.xpro.2021.100961.

Hensel, Jonathan A., Vinayak Khattar, Reading Ashton, and Selvarangan Ponnazhagan. 2019. “Characterization of Immune Cell Subtypes in Three Commonly Used Mouse Strains Reveals Gender and Strain-Specific Variations.” Laboratory Investigation 99 (1): 93–106. 10.1038/s41374-018-0137-1.

Herold, Kevan C., Thomas Delong, Ana Luisa Perdigoto, Noah Biru, Todd M. Brusko, and Lucy S.K. Walker. 2024. “The Immunology of Type 1 Diabetes.” Nature Reviews Immunology. Nature Research. 10.1038/s41577-023-00985-4.

Ju, Huai Qiang, Yun Xin Lu, Dong Liang Chen, Zhi Xiang Zuo, Ze Xian Liu, Qi Nian Wu, Hai Yu Mo, et al. 2019. “Modulation of Redox Homeostasis by Inhibition of MTHFD2 in Colorectal Cancer: Mechanisms and Therapeutic Implications.” Journal of the National Cancer Institute 111 (6): 584–96. 10.1093/jnci/djy160.

Kawai, Junya, Tadashi Toki, Masahiro Ota, Hidekazu Inoue, Yoshimi Takata, Takashi Asahi, Makoto Suzuki, et al. 2019. “Discovery of a Potent, Selective, and Orally Available MTHFD2 Inhibitor (DS18561882) with in Vivo Antitumor Activity.” Journal of Medicinal Chemistry. 10.1021/acs.jmedchem.9b01113.

Kim, Joungmok, Goowon Yang, Yeji Kim, Jin Kim, and Joohun Ha. 2016. “AMPK Activators: Mechanisms of Action and Physiological Activities.” Experimental and Molecular Medicine. Nature Publishing Group. 10.1038/emm.2016.16.

Kong, Byung Soo, Se Hee Min, Changhan Lee, and Young Min Cho. 2021. “MitochondrialEncoded MOTS-c Prevents Pancreatic Islet Destruction in Autoimmune Diabetes.” Cell Reports 36 (4). 10.1016/j.celrep.2021.109447.

Kory, Nora, Gregory A. Wyant, Gyan Prakash, Jelmi uit De Bos, Francesca Bottanelli, Michael E. Pacold, Sze Ham Chan, et al. 2018. “SFXN1 Is a Mitochondrial Serine Transporter Required for One-Carbon Metabolism.” Science 362 (6416). 10.1126/science.aat9528.

Kurniawan, Henry, Takumi Kobayashi, and Dirk Brenner. 2021. “The Emerging Role of OneCarbon Metabolism in T Cells.” Current Opinion in Biotechnology. Elsevier Ltd. 10.1016/j.copbio.2020.12.001.

Lamarre, Simon G., Luke MacMillan, Gregory P. Morrow, Edward Randell, Theerawat Pongnopparat, Margaret E. Brosnan, and John T. Brosnan. 2014. “An Isotope-Dilution, GCMS Assay for Formate and Its Application to Human and Animal Metabolism.” Amino Acids 46 (8): 1885–91. 10.1007/s00726-014-1738-7.

Lamarre, Simon G., Gregory Morrow, Luke MacMillan, Margaret E. Brosnan, and John T. Brosnan. 2013. “Formate: An Essential Metabolite, a Biomarker, or More?” In Clinical Chemistry and Laboratory Medicine, 51:571–78. 10.1515/cclm-2012-0552.

Li, Yuchan, Omar Elakad, Sha Yao, Alexander von Hammerstein-Equord, Marc Hinterthaner, Bernhard C. Danner, Carmelo Ferrai, Philipp Ströbel, Stefan Küffer, and Hanibal Bohnenberger. 2022. “Regulation and Therapeutic Targeting of MTHFD2 and EZH2 in KRAS-Mutated Human Pulmonary Adenocarcinoma.” Metabolites 12 (7). 10.3390/metabo12070652.

Li, Yujing, Minglong Cai, Yi Qin, Xiaojuan Dai, Liyuan Liang, Zhenyu Li, Xi Wen, Huizhi Jin, Chao Yang, and Zhu Chen. 2025. “MTHFD2 Promotes Osteoclastogenesis and Bone Loss in Rheumatoid Arthritis by Enhancing CKMT1-Mediated Oxidative Phosphorylation.” BMC Medicine 23 (1). 10.1186/s12916-025-03945-y.

Lieberman, Scott M, Anne M Evans, Bingye Han, Toshiyuki Takaki, Yuliya Vinnitskaya, Jennifer A Caldwell, David V Serreze, et al. 2003. “Identification of the Cell Antigen Targeted by a Prevalent Population of Pathogenic CD8 T Cells in Autoimmune Diabetes.” www.pnas.orgcgidoi10.1073pnas.0932778100.

Linsley, Peter S., and S. Alice Long. 2019. “Enforcing the Checkpoints: Harnessing T-Cell Exhaustion for Therapy of T1D.” *Current Opinion in Endocrinology*, Diabetes and Obesity. Lippincott Williams and Wilkins. 10.1097/MED.0000000000000488.

Liu, Shuyu, Xiaoke Wang, Liye Zhao, Liangren Zhang, and Yali Song. 2025. “MTHFD2: A Significant Mitochondrial Metabolic Enzyme and a Novel Target for Anticancer Therapy.” Drug Discovery Today. Elsevier Ltd. 10.1016/j.drudis.2025.104372.

Luo, Lan, Yuhong Chen, Xiao Chen, Yongwei Zheng, Vivian Zhou, Mei Yu, Robert Burns, et al. 2020. “Kras-Deficient T Cells Attenuate Graft-versus-Host Disease but Retain Graft-versusLeukemia Activity.” The Journal of Immunology 205 (12): 3480–90. 10.4049/jimmunol.2000006.

Ma, Eric H., Glenn Bantug, Takla Griss, Stephanie Condotta, Radia M. Johnson, Bozena Samborska, Nello Mainolfi, et al. 2017. “Serine Is an Essential Metabolite for Effector T Cell Expansion.” Cell Metabolism 25 (2): 345–57. 10.1016/j.cmet.2016.12.011.

Marrero, Idania, Allen Vong, Yang Dai, and Joanna D. Davies. 2012. “T Cell Populations in the Pancreatic Lymph Node Naturally and Consistently Expand and Contract in NOD Mice as Disease Progresses.” Molecular Immunology 52 (1): 9–18. 10.1016/j.molimm.2012.04.004.

Nakayasu, Ernesto S., Lisa M. Bramer, Charles Ansong, Athena A. Schepmoes, Thomas L. Fillmore, Marina A. Gritsenko, Therese R. Clauss, et al. 2023. “Plasma Protein Biomarkers Predict the Development of Persistent Autoantibodies and Type 1 Diabetes 6 Months Prior to the Onset of Autoimmunity.” Cell Reports Medicine 4 (7). 10.1016/j.xcrm.2023.101093.

Nilsson, Roland, Mohit Jain, Nikhil Madhusudhan, Nina Gustafsson Sheppard, Laura Strittmatter, Caroline Kampf, Jenny Huang, Anna Asplund, and Vamsi K. Mootha. 2014. “Metabolic Enzyme Expression Highlights a Key Role for MTHFD2 and the Mitochondrial Folate Pathway in Cancer.” Nature Communications 5 (January). 10.1038/ncomms4128.

Ogle, Graham D., Steven James, Dana Dabelea, Catherine Pihoker, Jannet Svennson, Jayanthi Maniam, Emma L. Klatman, and Chris C. Patterson. 2022. “Global Estimates of Incidence of Type 1 Diabetes in Children and Adolescents: Results from the International Diabetes Federation Atlas, 10th Edition.” Diabetes Research and Clinical Practice 183 (January). 10.1016/j.diabres.2021.109083.

Pardo-Lorente, Natalia, and Sara Sdelci. 2024. “MTHFD2 in Healthy and Cancer Cells: Canonical and Non-Canonical Functions.” Npj Metabolic Health and Disease 2 (1). 10.1038/s44324-024-00005-6.

Pearson, James A., F. Susan Wong, and Li Wen. 2016. “The Importance of the Non Obese Diabetic (NOD) Mouse Model in Autoimmune Diabetes.” Journal of Autoimmunity. Academic Press. 10.1016/j.jaut.2015.08.019.

Pietzke, Matthias, Johannes Meiser, and Alexei Vazquez. 2020. “Formate Metabolism in Health and Disease.” Molecular Metabolism. Elsevier GmbH. 10.1016/j.molmet.2019.05.012.

Pollizzi, Kristen N., and Jonathan D. Powell. 2015. “Regulation of T Cells by MTOR: The Known Knowns and the Known Unknowns.” Trends in Immunology. Elsevier Ltd. 10.1016/j.it.2014.11.005.

Rao, Enyu, Yuwen Zhang, Qiang Li, Jiaqing Hao, Nejat K Egilmez, Jill Suttles, and Bing Li. 2016. “AMPK-Dependent and Independent Effects of AICAR and Compound C on T-Cell Responses.” www.impactjournals.com/oncotarget.

Ren, Xinxin, Xiang Wang, Guowan Zheng, Shanshan Wang, Qiyue Wang, Mengnan Yuan, Tong Xu, Jiajie Xu, Ping Huang, and Minghua Ge. 2024. “Targeting One-carbon Metabolism for Cancer Immunotherapy.” Clinical and Translational Medicine 14 (1). 10.1002/ctm2.1521.

Ron-Harel, Noga, Daniel Santos, Jonathan M. Ghergurovich, Peter T. Sage, Anita Reddy, Scott B. Lovitch, Noah Dephoure, et al. 2016. “Mitochondrial Biogenesis and Proteome Remodeling Promote One-Carbon Metabolism for T Cell Activation.” Cell Metabolism 24 (1): 104–17. 10.1016/j.cmet.2016.06.007.

Rowe, Jared H., Ilaria Elia, Osmaan Shahid, Emily F. Gaudiano, Natalia E. Sifnugel, Sheila Johnson, Amy G. Reynolds, et al. 2023. “Formate Supplementation Enhances Antitumor CD8+ T-Cell Fitness and Efficacy of PD-1 Blockade.” Cancer Discovery 13 (12): 2566–83. 10.1158/2159-8290.CD-22-1301.

Saleiro, Diana, and Leonidas C. Platanias. 2015. “Intersection of MTOR and STAT Signaling in Immunity.” Trends in Immunology. Elsevier Ltd. 10.1016/j.it.2014.10.006.

Shang, Man, Lina Ni, Xiao Shan, Yan Cui, Penghui Hu, Zemin Ji, Long Shen, et al. 2023. “MTHFD2 Reprograms Macrophage Polarization by Inhibiting PTEN.” Cell Reports 42 (5). 10.1016/j.celrep.2023.112481.

Shieh, J. J., C. J. Pan, B. C. Mansfield, and J. Y. Chou. 2005. “In Islet-Specific Glucose-6Phosphatase-Related Protein, the Beta Cell Antigenic Sequence That Is Targeted in Diabetes Is Not Responsible for the Loss of Phosphohydrolase Activity.” Diabetologia 48 (9): 1851–59. 10.1007/s00125-005-1848-6.

Sugiura, Ayaka, Gabriela Andrejeva, Kelsey Voss, Darren R. Heintzman, Xincheng Xu, Matthew Z. Madden, Xiang Ye, et al. 2022. “MTHFD2 Is a Metabolic Checkpoint Controlling Effector and Regulatory T Cell Fate and Function.” Immunity 55 (1): 65–81.e9. 10.1016/j.immuni.2021.10.011.

Tang, Hui, and Ning Hou. 2023. “Whether MTHFD2 Plays a New Role: From Anticancer Targets to Anti-Inflammatory Disease.” Frontiers in Pharmacology 14. 10.3389/fphar.2023.1257107.

Thompson, Peter J., Ajit Shah, Vasilis Ntranos, Frederic Van Gool, Mark Atkinson, and Anil Bhushan. 2019. “Targeted Elimination of Senescent Beta Cells Prevents Type 1 Diabetes.” Cell Metabolism 29 (5): 1045–1060.e10. 10.1016/j.cmet.2019.01.021.

Tonnerre, Pierre, David Wolski, Sonu Subudhi, Jihad Aljabban, Ruben C. Hoogeveen, Marcos Damasio, Hannah K. Drescher, et al. 2021. “Differentiation of Exhausted CD8+ T Cells after Termination of Chronic Antigen Stimulation Stops Short of Achieving Functional T Cell Memory.” Nature Immunology 22 (8): 1030–41. 10.1038/s41590-021-00982-6.

Vitto, Humberto De, Danushka B. Arachchige, Brian C. Richardson, and Jarrod B. French. 2021. “The Intersection of Purine and Mitochondrial Metabolism in Cancer.” Cells. MDPI. 10.3390/cells10102603.

Wiedeman, Alice E., Virginia S. Muir, Mario G. Rosasco, Hannah A. DeBerg, Scott Presnell, Bertrand Haas, Matthew J. Dufort, et al. 2020. “Autoreactive CD8+ T Cell Exhaustion Distinguishes Subjects with Slow Type 1 Diabetes Progression.” Journal of Clinical Investigation 130 (1): 480–90. 10.1172/JCI126595.

Yang, Kangping, Yihan Zhang, Jiatong Ding, Zelin Li, Hejin Zhang, and Fang Zou. 2024. “Autoimmune CD8+ T Cells in Type 1 Diabetes: From Single-Cell RNA Sequencing to TCell Receptor Redirection.” Frontiers in Endocrinology. Frontiers Media SA. 10.3389/fendo.2024.1377322.

Zarou, Martha M., Alexei Vazquez, and G. Vignir Helgason. 2021. “Folate Metabolism: A ReEmerging Therapeutic Target in Haematological Cancers.” Leukemia. Springer Nature. 10.1038/s41375-021-01189-2.

Zhan, Jun Kun, Yan Jiao Wang, Shuang Li, Yi Wang, Pan Tan, Je Yu He, Yi Yin Chen, et al. 2018. “AMPK/TSC2/MTOR Pathway Regulates Replicative Senescence of Human Vascular Smooth Muscle Cells.” Experimental and Therapeutic Medicine 16 (6): 4853–58. 10.3892/etm.2018.6767.

Zhang, Jiaxue, Tong Lyu, Yaming Cao, and Hui Feng. 2021. “Role of TCF-1 in Differentiation, Exhaustion, and Memory of CD8+ T Cells: A Review.” FASEB Journal. John Wiley and Sons Inc. 10.1096/fj.202002566R.

Zhao, Xudong, Qiang Shan, and Hai Hui Xue. 2022. “TCF1 in T Cell Immunity: A Broadened Frontier.” Nature Reviews Immunology. Nature Research. 10.1038/s41577021-00563-6.

Zhou, Xin, Yanghao Zhong, Olivia Molinar-Inglis, Maya T. Kunkel, Mingyuan Chen, Tengqian Sun, Jiao Zhang, et al. 2020. “Location-Specific Inhibition of Akt Reveals Regulation of MTORC1 Activity in the Nucleus.” Nature Communications 11 (1). 10.1038/s41467-020-19937-w.

Zhu, Zhiyuan, and Gilberto Ka Kit Leung. 2020. “More Than a Metabolic Enzyme: MTHFD2 as a Novel Target for Anticancer Therapy?” Frontiers in Oncology. Frontiers Media S.A. 10.3389/fonc.2020.00658.

